# Phenotypic AI-based design of cell-specific small molecule cytotoxics

**DOI:** 10.1101/2025.10.15.682546

**Authors:** Gema Rojas-Granado, Marta Sánchez-Soto, Jesús Calahorra, María Caballero, Israel Ramos, Martino Bertoni, Patrick Aloy

## Abstract

Drug discovery is being transformed by artificial intelligence, which enables the exploration of vast chemical spaces and the generation of novel compounds with tailored properties. While most generative approaches focus on optimizing physicochemical features or protein-binding affinities, their application to phenotypic drug discovery remains limited. Here we present an integrated framework to design small molecules with selective cytotoxic effects in pancreatic cancer cells that combines high-throughput cytotoxicity screening, bioactivity signature–based machine learning predictors, and reinforcement learning–driven generative models. We screened over 11,000 compounds across six pancreatic cancer and two control cell lines, confirming 392 hits, and used these data to train cell line–specific classifiers that outperformed fingerprint-based models. We embedded these predictors into the REINVENT platform to generate molecules with desired cytotoxicity profiles, including challenging cases of distinguishing between pancreatic cancer lines with highly similar molecular backgrounds. We generated between 37 and 137 high-scoring candidates in seven different cell selectivity exercises, which were structurally distant from the active compounds in the training set. We tested 45 close analogues to the AI-designed molecules, and 20 of them showed the desired cell-specific cytotoxic effects. Indeed, we validated at least one compound in 5 of the 7 exercises tested, showing over a large increase in hit discovery rate compared to the HTS. Additionally, we benchmarked our strategy against direct predictions on a library of over 150,000 compounds, skipping the generative step, finding that the fraction of hits, and more importantly the dose-response validation, is markedly higher in the AI-designed molecules. Overall, our study demonstrates the feasibility of combining predictive and generative AI methods to design molecules with complex phenotypic outcomes in a target-agnostic manner, paving the way for next-generation phenotype-driven drug discovery strategies.

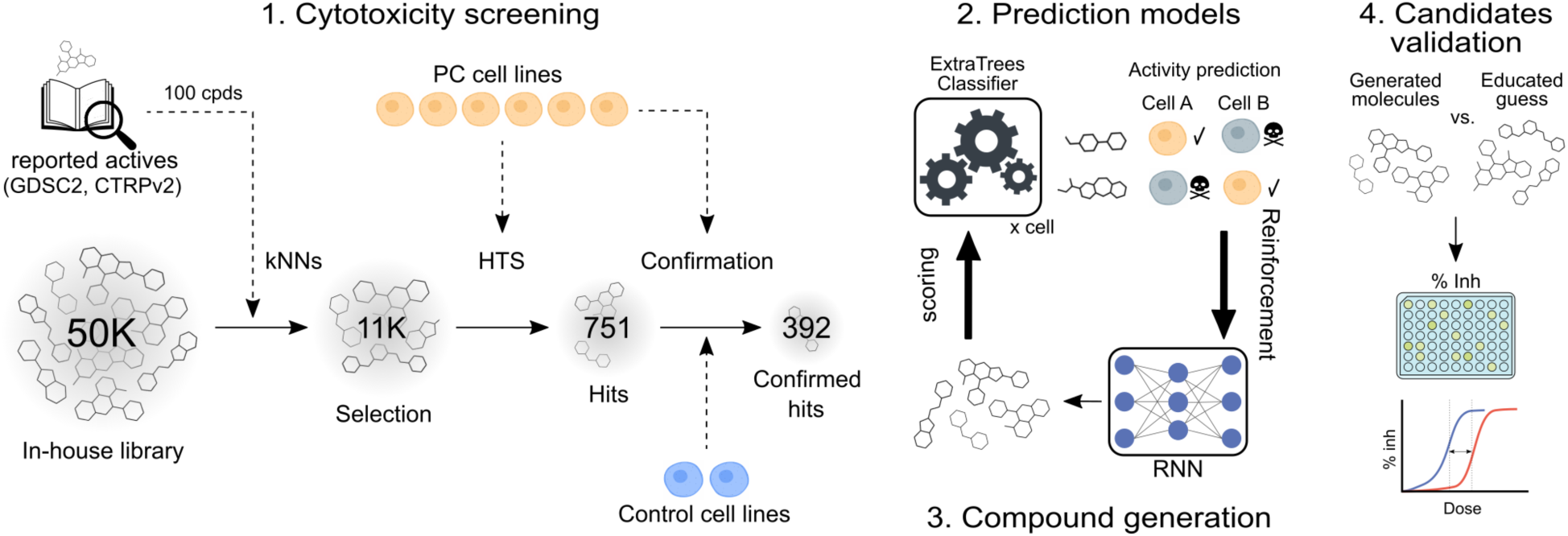

## Main

Drug discovery begins with identifying small-molecule hit compounds that often exhibit low-affinity interactions with biological targets, which usually undergo chemical optimization to enhance affinity, specificity, and drug-like properties. Identifying promising hits early is critical for downstream success, and high-throughput screenings (HTS) allow rapid evaluation of compound libraries^1^. As chemical collections have expanded exponentially, so too have the opportunities to identify promising drug candidates. Historically, pharmaceutical companies maintained compound collections containing millions of molecules, but advances in combinatorial chemistry, ultra-large chemical synthesis, and artificial intelligence-driven *de novo* molecule design have propelled library sizes into the billions and even trillions^2^. Ultra-large compound libraries offer unparalleled chemical diversity, significantly improving the odds of finding novel, high-affinity interactions with therapeutic targets. However, this growth has introduced new challenges in computational drug discovery, as traditional molecular docking methods struggle to efficiently process such immense chemical spaces^3^. To address this scalability issue, new computational strategies are being developed that leverage machine learning (ML) and chemical descriptors to efficiently navigate ultra-large libraries^4–7^. These approaches aim to pre-filter vast chemical spaces, identifying the most promising subsets of compounds for subsequent docking and experimental validation.

An alternative to the exploration of ultra-large compound libraries comes from generative artificial intelligence (genAI), which integrates AI-driven molecular property prediction and generative modeling. Indeed, while virtual screening mostly focuses on assessing the potential binding of existing compounds to specific targets, deep generative models have shifted the paradigm by enabling the creation of novel molecular structures. For instance, generative adversarial networks (GANs) have proved effective at discovering DDR1 kinase inhibitors in just 21 days, revolutionizing the traditional drug discovery timeline^8^, and chemical language models (CLM) have generated molecules with specific dual-protein binding profiles^9^, showcasing the potential for target and multi-target -based drug design. Additionally, reinforcement learning (RL)-based platforms, like REINVENT^10,11^, have further advanced this field by allowing molecular generators to optimize multiple property objectives in an easy-to-use, modular framework compatible with traditional QSAR models. More specifically, recent developments have focused on the generation of molecules with particular physicochemical properties or protein-binding capabilities^8,9,12,13^.

Postgenomic initiatives have identified many novel potential pharmacological targets^14,15^, offering a wealth of opportunities for therapeutic intervention. However, the deluge of molecular and cellular data collected in the last years has also shown that, given the robustness of biological systems, the modulation of a single target is seldom exquisite, highlighting the limitations of target-based drug discovery approaches. Recent analyses have shown that phenotypic screens, which focus on cellular or physiological outcomes without prior assumptions about the mechanism, are behind most of the FDA-approved drugs available nowadays, and only 9.4% of small molecule drugs have been discovered through target-based assays^16^. Indeed, the more general phenotype-based approaches seem to better capture the complex, multi-pathway, interactions that drive cellular responses, contributing to the discovery of first-in-class drugs, while target-centric approaches appear more useful for the development of follow-on compounds^17,18^.

In an attempt to seize biological complexity, it is increasingly common for high-throughput experiments to simultaneously characterize multiple omics profiles (i.e. trans-omics analyses)^19,20^, so that several views of a given phenomenon can be analysed simultaneously. This strategy is also applied to the study of drug-induced perturbations, with several studies measuring, for instance, the global effects caused by small molecules in the transcriptome^21^, proteome^22^ or metabolome^23^ of different cell types. In parallel, numerous methodologies to integrate these diverse data are emerging, with the aim of capturing the coordinated interplay of the many regulatory layers present in biological systems^24^. We recently developed the Bioteque, a resource of unprecedented size and scope that contains pre-calculated biomedical descriptors derived from a gigantic knowledge graph, displaying more than 450 thousand biological entities and 30 million relationships between them^25,26^. Overall, the Bioteque contains over 1,000 signatures to capture (embed) the biological context of different biological entities (genes, cells, tissues, diseases, etc.), which facilitate understanding of how cells work at a multi-scale resolution. From the perspective of the drugs, we also profited from embedding strategies to integrate the major chemogenomics and drug activity repositories in a single resource named the Chemical Checker (CC), which represents the largest collection of small molecule bioactivity signatures available to date^27^. The CC divides data into five levels of increasing complexity, ranging from the chemical properties of compounds to their clinical outcomes. To overcome the scarcity of experimental bioactivity data, we implemented a collection of deep neural networks (i.e. *Signaturizers*) able to infer bioactivity signatures for any compound of interest, even when little or no experimental information is available for them^28,29^. The use of bioactivity signatures has facilitated, for instance, the identification of compounds that elicit the same phenotypic effects than monoclonal antibodies used in clinical practice, and small molecules to indirectly target SNAIL1, a transcription factor involved in EMT processes linked to colorectal cancer and classified as undruggable^27,28^. Overall, the embedding of biological and bioactivity data associated to small molecules opens new opportunities for computational drug discovery. On the one hand, the vector-like format of these descriptors is the ideal input for most machine learning algorithms and, on the other, placing biology and chemistry in the same mathematical space they facilitate the identification of connections between chemical compounds and the effects that they trigger when used on biological samples. Effectively, this means that specific bioactivity properties of the input molecules can be learned together with their chemical structure.

Despite the growing interest in deep learning-based *de novo* molecular design, the application of generative models to phenotypic drug discovery remains largely unexplored, partly because the lack of specific training data. Indeed, most generative efforts to date have focused on optimizing physicochemical properties or designing compounds that bind to specific protein targets^8–13^. There are, however, some remarkable exceptions that show the feasibility of the approach. For instance, Joo et al.^30^ first applied a conditional variational autoencoder to generate molecular fingerprints that are predicted to inhibit cell growth. Similarly, Mendez-Lucio et al.^31^, used genAI models to design molecules able to induce specific transcriptional responses in cells, bridging the gap between chemical structures and cellular gene expression profiles. Building on these concepts, subsequent studies have generated compounds aimed at inducing desired transcriptional states^32–34^, and more recently, Kim et al.^35^ applied a similar strategy to design cytotoxic agents tailored to specific genotypic profiles. Yet, these efforts remain confined to the computational domain, as experimental validation of the proposed candidates is still lacking, underscoring the challenge of translating such models into biological systems.

In this study, we explore the application of genAI to design small molecule cytotoxics that are cell-selective across multiple pancreatic cancer cell lines. In brief, we first run a high-throughput compound screen to build the dataset to train our cytotoxicity prediction models. We next develop and validate classifiers based on bioactivity signatures, and couple them with a generative framework to design new compounds for each desired cytotoxicity profile. Finally, we experimentally validate several of these generated compounds, confirming that genAI can indeed be harnessed to create novel chemical entities exhibiting the desired cell-selectivity profiles without relying on predefined molecular targets.

## Results

### Collecting cell line sensitivity data

To set the cellular framework of our experiment, we selected a panel of well-characterised pancreatic cancer (PC) cells comprising six lines with increasing levels of mutation: from CAPAN2, which has mutations only in *KRAS*, to BXPC3, with four different mutations, none of them in the *KRAS* gene (*CDKN2A*, *MAP2K4*, *SMAD4*, and *TP53*). On top of their distinct mutational profiles, they also differ in origin of the source tumour (i.e. primary tumour or metastases at different locations) and histology (adenocarcinoma or ductal adenocarcinoma) (Supplementary Table 1). Overall, despite some differences, these cells show quite similar transcriptional profiles, and they can be clearly differentiated from cell lines derived from other tissues (Extended Data Fig.1). Finally, in addition to the pancreatic cancer cells, we included two non-pancreatic lines to evaluate our approach across different cell types: the osteosarcoma U2OS and the non-tumorigenic HEK293, both of which are widely used in research.

We first retrieved information on small molecules cytotoxics active on the pancreatic cancer cell lines available in the public domain, to evaluate its suitability to build accurate cell line-specific sensitivity prediction models in our panel. Supplementary Table 2 summarises the different resources we examined and the number of drugs and cell lines tested in each. In brief, NCI-60^36^ is the largest resource, with more than 20,000 drugs screened across 60 human tumour cell lines; however, it does not include any pancreatic cancer cell lines. By contrast, CTRPv2^37^ and GDSC2^38^ contain approximately 200 and 450 different drugs, respectively, tested on more than 800 cell lines, including all the pancreatic ones we selected for this study. Upon closer examination of these two resources, we found 237 molecules active on any of our target cell lines out of the 583 tested (Extended Data Fig. 2a). For the 38 compounds present in both resources, we checked the robustness of the reported cytotoxicities, and we found a poor correlation in three of the six PC cell lines, although we could not identify any highly active compound present in both (Extended Data Fig. 2a, scatter plots). Additionally, to further evaluate the utility of this data within our project, we selected 12 drugs showing differential activity for the PC cell lines from DrugCell^39^ (a resource integrating the data from CTRPv2 and GDSC), and tested them in the lab to see whether we could reproduce the reported results (Supplementary Table 3). Extended Data Fig. 2b shows how only 3 of the 12 drugs behaved as expected when assessed through a dose-response MTT assay. These discouraging findings, combined with the lack of cytotoxicity data for non-tumorigenic cell lines and previous concerns in the community regarding reproducibility^40,41^, led us to generate a larger, more reliable in-house dataset for cell line sensitivity to train our models.

We performed a high-throughput screening (HTS) with more than 11,000 molecules across all the pancreatic cancer cell lines included in our panel (Fig. 1a). We used the IRB in-house library of compounds which, at that time, comprised over 47,000 different molecules from the ChemDiv screening libraries. To run our HTS, we selected 35 plates containing 11,053 molecules. To increase our chances of identifying active compounds with diverse specificity, we first selected 100 seed compounds from DrugCell that exhibited different cytotoxicity profiles across the pancreatic cancer and U2OS cells. We then used small molecule bioactivity descriptors from the CC to identify similar compounds in the IRB library, and selected the 35 plates with the highest number of analogous similar to the seed ones (see *Methods*). Fig. 1c shows a representation of the chemical space covered by the compounds tested in this HTS compared to that covered by the NCI-60, CTRPv2 and GDSC2 datasets. As expected, we found a significant enrichment of hit compounds within the set of molecules leading the plate selection (Supplementary Table 4).

**Fig. 1.**
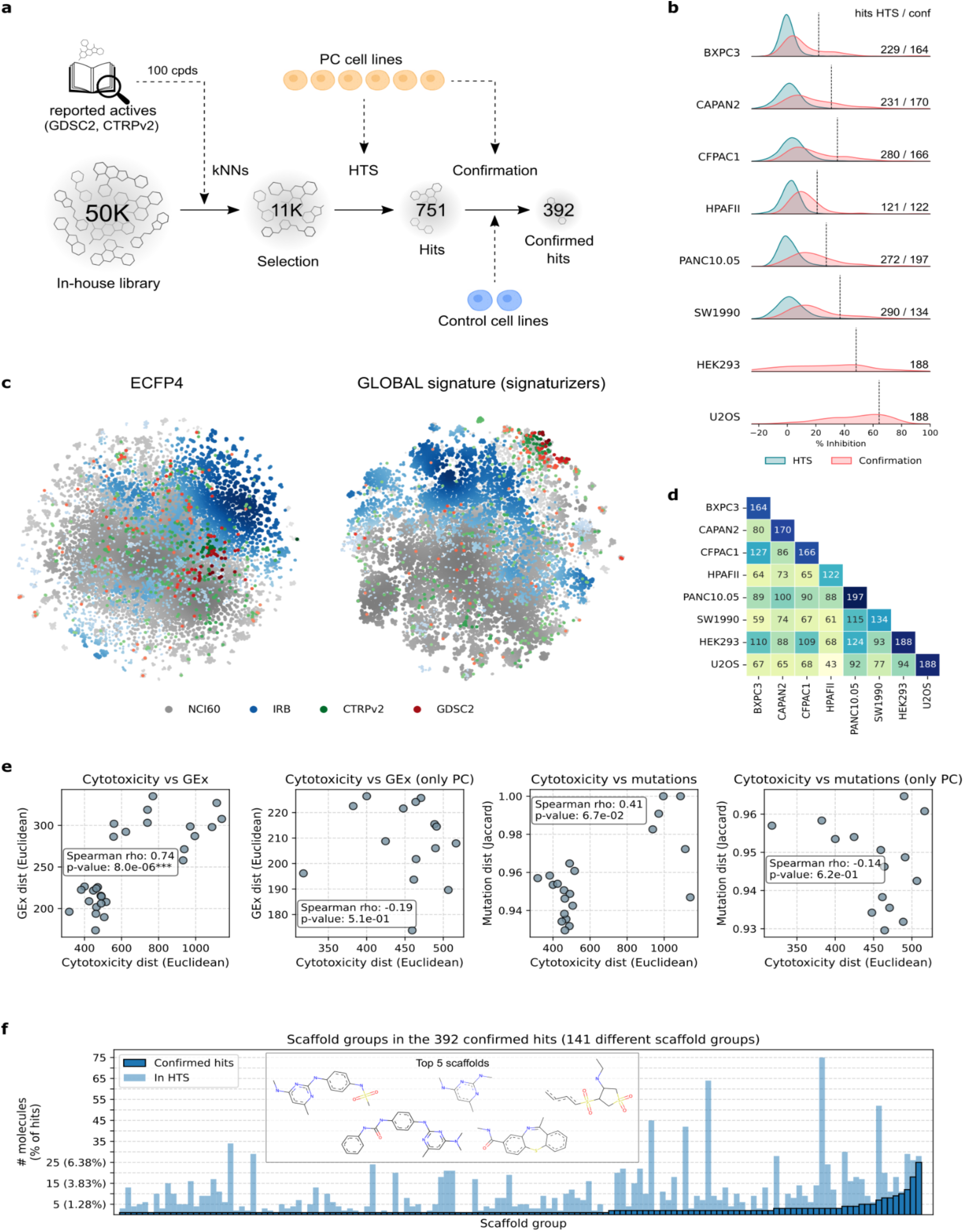
High Throughput Screening (HTS) methodology and results. **(a)** A total of 35 plates comprising 11,053 compounds were selected, prioritizing those enriched with molecules similar to reported actives in the literature. The compounds were screened on a panel of six pancreatic cancer (PC) cell lines, and the hits were confirmed in a second experiment that included two control non-pancreatic cell lines. **(b)** Distribution of activity (% inhibition) for the compounds tested in the HTS (blue) and confirmation (red) experiments across cell lines, and number of hits. Dashed vertical lines indicate sensitivity thresholds. **(c)** tSNE (t-distributed Stochastic Neighbor Embedding) representation of molecules from different drug sensitivity databases (GDSC2, CTRPv2 and NCI-60) and the in-house library (IRB). Each molecule is represented as a single dot, colored and sized by 2D density within the respective dataset. ECFP4 corresponds to Extended Connectivity Fingerprints molecular representations^43^, while GLOBAL signatures are molecular bioactivity descriptors coming from the *Signaturizers*^28^ (see *Methods*). **(d)** Overlap of hits between cell line pairs in the confirmation experiment. The diagonal represents number of hits per cell line. **(e)** Correlation between cell line distances in cytotoxicity (from the confirmation experiment) and gene expression (GEx) or mutational profiles (see *Methods*). The first and third plots include all cell lines (six pancreatic plus controls), while the second and fourth focus on PC cells only. **(f)** Molecular scaffolds present in active molecules from the confirmation experiment, showing how many active molecules contain each scaffold (dark blue). Lighter bars represent the total number of molecules with those scaffolds in the HTS. Differences between the bars for a given scaffold reflect inactive molecules. The top 5 scaffold groups are shown.

Next, we tested the 11,000 molecules on the 6 pancreatic cancer cell lines at 10 µM, obtaining a total of 751 hits, ranging from 121 in HPAFII to 290 in SW1990 cells. Given the potentially different sensitivities displayed by each cell line, we defined the hits as those molecules exhibiting a percentage inhibition higher than the mean plus 2.5 standard deviations, calculated from the entire distribution of 11,000 compounds for each individual cell line (see *Methods*). Thus, activity thresholds are cell-line dependent, ranging from a death rate of 20.9% in HPAFII to 36.9% in SW1990, where lines that are generally more difficult to kill show lower thresholds than those that are more sensitive overall (Fig. 1b). We then confirmed the hits in a second experiment, where we re-tested the 751 hit molecules at 10 µM, this time including the auxiliary cell lines U2OS and HEK293. Overall, we confirmed 392 hits in the PC panel, ranging from 122 in HPAFII to 197 in PANC10.05 cells (Fig. 1b; Supplementary Table 5). Please, note that some HTS non-active borderline compounds in one cell-line might be hits in another, and thus re-tested for confirmation in all cell lines, which could result in a higher number of confirmed hits per line. We observed the auxiliary cell lines to be much more sensitive to drug treatment than any of the pancreatic cancer ones, particularly in the case of the U2OS cells, where the median inhibition exceeded 50%. For those 2 cell lines, since we did not have a background distribution to use as a reference, we defined the activity thresholds as the 75th percentile of all the compounds tested, which resulted in a similar number of active molecules to those found in the other cells (188 for each). Fig. 1d shows the overlap in hit molecules for each pair of cell lines following this binarization criteria.

We then explored the potential relationship between the sensitivity profiles and the biological background of the cells, specifically their mutational and gene expression patterns. A naïve conjecture would suggest that cell lines with more similar mutational or gene expression profiles would also show greater similarity in terms of sensitivity. To test this hypothesis, we built vector-like compound sensitivity profiles per each cell line, including the results for the initial 751 hit compounds in the confirmation experiments, and computed Euclidean distances between profile pairs. We also downloaded mutational profiles for each cell line from DrugCell. These are 3,008- dimensional vectors where each dimension represents a gene and 1 or 0 denotes whether it is mutated or not, respectively. We then filtered out the genes that were not mutated in any of the cell lines and ended up with 291-dimensional mutational descriptors. In this case, since we are dealing with binary fingerprints, we computed Jaccard’s distances between pairs of cells to find mutational distances. Finally, we conducted RNA-seq experiments to examine the transcriptional signatures of each cell line, and used Euclidean distances between pairs for the posterior analysis. In this case, we did not rely on published profiles to avoid artefacts due to potential drifts in cell cultures (Supplementary Table 6). To compare the three different cellular profiles (i.e. drug sensitivity, mutations and gene expression), we computed Spearman’s correlations between the distances found by each approach (Fig. 1e). We observed moderate to high correlations between the distances in both gene expression and mutations with the distances in sensitivity profiles when considering the entire cell line panel (rho=0.74, p-value<0.001 for sensitivity vs gene expression; rho=0.41, p-value=0.07 for sensitivity vs mutation). However, when we removed the non-pancreatic cell lines from the comparisons, the correlations disappeared. In other words, pancreatic cancer cell lines with similar mutational or gene expression profiles show different sensitivity to drug-like molecules (Spearmańs correlations of -0.19 for sensitivity vs gene expression, -0.14 for sensitivity vs mutation, with p-values>0.5). We also explored possible correlations with proteomic profiles from DepMap^42^ and Bioteque descriptors^25,26^, capturing complex relationships between different cellular entities (Extended Data Fig. 3). Overall, we observed little or no correlations between drug sensitivity profiles and any of the descriptors explored when considering only pancreatic cancer cell lines, since they are probably too similar to reveal direct correlations. However, we indeed find clear differences between pancreatic cancer and the auxiliary cell lines U2OS and HEK293.

Finally, we analyzed the chemical structure of the hit compounds to check for commonalities that could be indicative of scaffold-specific cytotoxic effects. We studied the annotated scaffold groups in our in-house library and we looked for structural groups enriched in active molecules. Our compounds library is chemically diverse by design, with more than 1,700 scaffold groups present in the over 11,000 molecules tested in the HTS. Each scaffold group contains on average 5 molecules. Only 254 groups include 10 or more compounds, with the largest group holding 77 compounds (0.7% of the total), and 531 groups containing only 1 molecule. Among the 392 active molecules in pancreatic cancer cell lines, we identified 141 scaffold groups, where 11 groups hosted more than 5 active compounds (Fig. 1f). The ratio of active molecules in these 11 groups was higher than the observed in the complete HTS, with odd ratios ranging from 2.48 to 160.66, and p-values <0.05 (Extended Data Fig. 4a). Of particular interest is scaffold group CL4745A, for which we tested 5 different molecules, all of which were active exclusively in CAPAN2 and not in the other cell lines (Extended Data Fig. 4).

### Cytotoxicity prediction models

We benefited from the rich cytotoxicity data generated to train computational models to predict whether each pancreatic cancer cell line would be sensitive or resistant to a given compound, so that we could later on use these models to bias the generation of new molecules. In particular, we trained ExtraTrees classifiers using all the identified active compounds for each cell line, and added 10 times as many inactive molecules, maintaining a fixed positive:negative ratio of 1:10 in the training dataset. To represent the compounds, we used either ECFP4 fingerprints^43^ or CC bioactivity signatures^28^ (CC_GLOBAL; Fig. 2a). See *Methods* for further details on the molecular descriptors and model building. We then evaluated the models *in silico* using a 10-fold cross-validation approach, and compared the performance of the two different molecular descriptors.

**Fig. 2.**
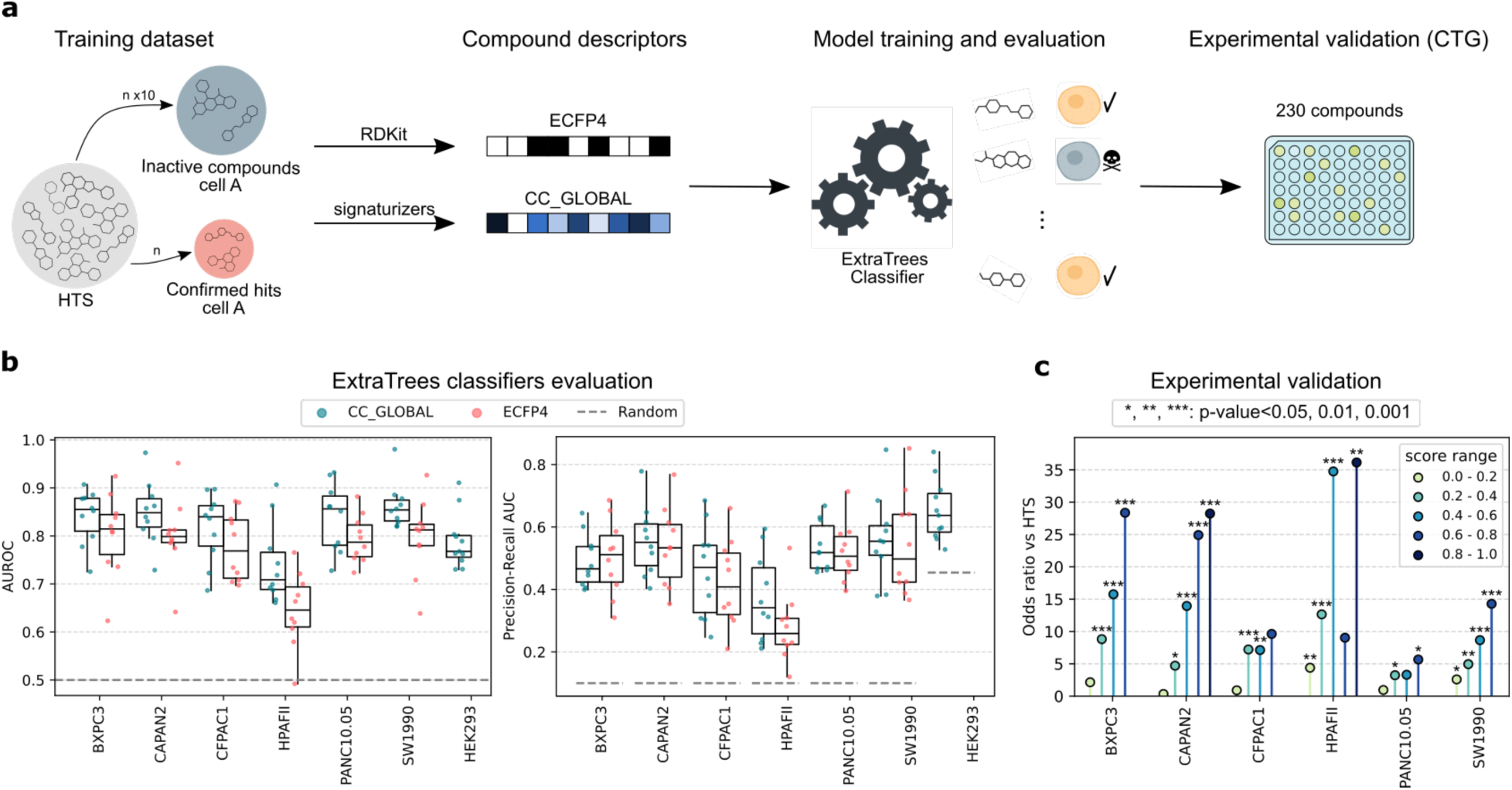
Building and evaluation of cytotoxicity prediction models. **(a)** To build cytotoxicity prediction models for each cell line, we selected all hits from the confirmation experiment and included ten times more inactive molecules. Compounds in the training set were represented as numerical vectors using ECFP4 or GLOBAL signature (CC_GLOBAL) descriptors (see *Methods*). ExtraTrees classifiers were trained individually for each cell line (six PC cells plus HEK293). The models were experimentally validated using a CellTiter-Glo survival assay (see Methods) with 230 compounds selected from the in-house library based on predicted scores (see *Methods*). **(b)** *In silico* evaluation of the cytotoxicity prediction models for each descriptor type (ECFP4 and CC_GLOBAL), using a random split, 10-fold cross-validation strategy. Left: area under the Receiver Operating Characteristic Curve (AUROC). Right: area under the precision-recall curve. Dotted lines indicate random expectation. **(c)** Experimental validation of the prediction models. A set of 230 compounds were selected across various ranges of predicted scores and Tanimoto similarity to previously tested molecules (see *Methods*). These compounds were tested on the whole cell line panel. The plot shows the enrichment of hits at different score ranges relative to the HTS, representing how many times more hits were found compared to a quasi-random selection (HTS). Statistical p-values come from Fisher’s exact test.

We observed that the models trained with small molecule bioactivity descriptors (CC_GLOBAL) consistently outperformed those trained with ECFP4 fingerprints, with all models showing mean AUROCs (Area Under the Receiver Operating Characteristic Curve) exceeding 0.8 and mean AUPRs (Area Under the Precision-Recall Curve) approaching or surpassing 0.5 (random expectation = 0.1) across folds (Fig. 2b). The HPAFII cell line was the only exception, showing lower performance metrics (mean AUC = 0.74, mean AUPR = 0.37), but still outperforming ECFP4 fingerprints (mean AUC = 0.64, mean AUPR = 0.28).

After the cross-validation, we used the complete set of positives as training, without a validation split, to train a final model for each cell line using CC bioactivity descriptors, as they showed superior performance. We then sought to prospectively assess the performance of the final models in a real-world scenario, evaluating not only their capacity to identify active compounds but also a possible interpretation of the model scores. We thus ran our drug sensitivity models on the ∼35,000 IRB library compounds that had not been used in the initial HTS. We grouped the molecules into four categories based on their Tanimoto index to their closest compound in the training dataset (<0.4, 0.4–0.6, 0.6–0.8, >0.8), and selected the top seven scoring compounds from each range in each cell line, up to a total of 320 compounds (Extended Data Fig. 5). The predicted scores for the selected compounds ranged from 0 to 0.93, while their Tanimoto index to any compound in the training set ranged from 0.23 to 0.98. Finally, we experimentally tested the cytotoxic activity of the selected molecules in each cell line at 10 µM, and measured the enrichment in the number of hits compared to the HTS (Supplementary Table 7).

Fig. 2c shows the results of the experimental validation. For all cell lines, we observed higher hit enrichments as the predicted scores for the tested compounds increased. Indeed, we observed strong enrichments when the predicted score exceeded 0.6 or 0.8, ranging from 5.7 in PANC10.05 to 36-fold in HPAFII cells, showing that high scores do indeed translate into higher hit rates. We found no strong correlation between the prediction score and the growth inhibition exhibited by the compounds (Extended Data Fig. 6). This was expected, since our models were only trained using binary data (i.e. active or not-active) for each compound at 10 µM. Overall, we found that the best performing cytotoxicity models were those for BXPC3 and CAPAN2 cells, with hit enrichments for the high scores reaching 30-fold. The SW1990 model also showed a good performance, with a clear correlation between scores and hit enrichments, and CFPAC1 predictors proved to be particularly effective in identifying inactive compounds. The validation of the HPAFII model showed strong hit enrichment for scores higher than 0.8, although we also found significant enrichments (i.e. 10-fold) for low score values. A closer inspection of the % inhibition data, showed that very few molecules were actually active, which lowered our confidence in the predictors for this cell line. PANC10.05 models also showed modest performances in the prospective real-world validation, with most tested molecules being inactive independently of the predicted score. Regarding the non-pancreatic cell lines, we built prediction models for HEK293 using the same process as for the rest (i.e. training with all the hits and CC bioactivity descriptors). However, in this case, we could not maintain the 1:10 positive:negative ratio due to the lack of inactive compounds for this cell line, which was only included in the hit validation phase. Instead, we adopted the 1:4 ratio established in the confirmation assay. Although we also observed an increase in the fraction of hits for higher prediction scores, with 7% of molecules being active for scores <0.2 vs 33% for scores >0.8, we could not calculate enrichments due to the lack of the baseline provided by the HTS experiment. We excluded U2OS from the analysis due to its high number of hits, with a sensitivity threshold exceeding 60% inhibition (Fig. 1b), making it unsuitable for the selective cytotoxicity exercise.

### Generation and evaluation of new selective small-molecule cytotoxics

After the assessment and experimental validation of our cell-specific cytotoxicity models, we selected the best performing predictors to guide the generation of new molecules towards diverse cell selectivity profiles. More specifically, we defined 7 distinct showcase exercises, each aiming to exert cytotoxicity in one cell line without affecting another. First, we targeted BXPC3 while sparing CAPAN2, as these two models demonstrated an excellent performance during experimental validation. Given the strong ability of the CFPAC1 model to identify inactive compounds, we designed a second exercise to kill CAPAN2 while sparing CFPAC1. Additionally, we sought to generate molecules that could selectively target a pancreatic cancer cell line without affecting non-tumorigenic cells, a relevant task from a therapeutic perspective. For this, we generated molecules aimed at killing BXPC3 or CAPAN2 while sparing HEK293. Lastly, as a purely academic exercise, we tested what we expected to be a simpler task: selectively killing a non-tumorigenic cell without affecting the pancreatic cancer cell lines. To this end, we designed three exercises targeting HEK293 while sparing BXPC3, CAPAN2, or CFPAC1, respectively.

As generative tool, we used REINVENT v3.2^10^, a modelling framework for *de novo* molecular design that combines SMILES-based recurrent neural networks (RNNs) with reinforcement learning to optimize molecules for specific objectives. More specifically, it consists of an agent that generates chemical structures and a scoring function that evaluates them, connected through a reinforcement learning loop that guides molecular optimization. We thus incorporated our cell-specific cytotoxic models as scoring functions in the REINVENT framework, and ran it for up to 2,000 reinforcement cycles, evaluating the generated molecules using the two cell models relevant to each exercise (Fig. 3a). We optimized the generation of the molecules simultaneously for the two properties (i.e. one line sensitive and the other resistant). However, this approach did not directly yield satisfactory mean scores for both objectives, and we thus applied a curriculum learning strategy. This is, we initially optimized the generation of molecules for one objective (i.e. killing one cell line) and subsequently we added the second constraint (i.e. sparing another cell line) once the first objective reached a mean score above 0.4. Additionally, for two exercises (BXPC3 vs CAPAN2 and BXPC3 vs HEK293) we found a common substructure present in all the proposed molecules (Extended Data Fig. 7). In these cases, we executed an extra REINVENT run penalizing structures containing this motif. Further details on the parametrization of the training strategy are available in *Methods* and Supplementary Table 8. After training the generative model, we sampled approximately 10,000 molecules from the optimized agent for downstream evaluation.

**Fig. 3.**
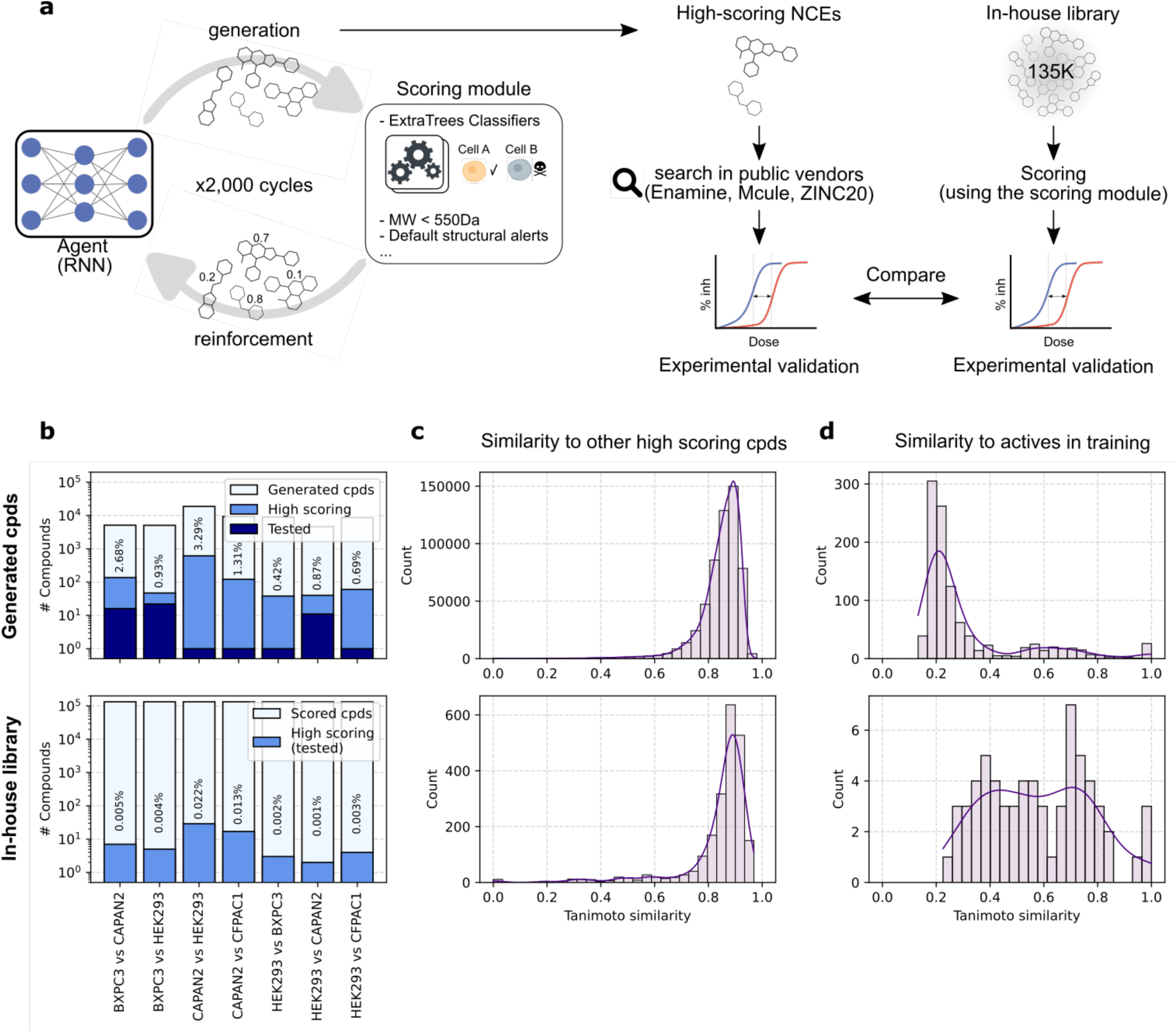
Generation of novel, cell line-specific cytotoxics. **(a)** We used REINVENT3.2^10^ as the generative framework to design new candidate molecules tailored to a specific cytotoxicity profile. We incorporated two cytotoxicity predictors in the scoring module: one for the target cell line and another to avoid affecting a second cell line. Additional filters, such as molecular weight and default structural alerts, were also included. We trained the agent for 2,000 steps of the reinforcement loop, and sampled 10,000 molecules from the trained agent. High-scoring molecules or their close analogs were searched in public vendor databases (Enamine, Mcule, and ZINC20^44–46^) and experimentally tested if found. In parallel, we made predictions on the extended in-house library (135,000 compounds), and tested the high scoring candidates for comparison. **(b)** Number of unique generated, high-scoring and tested compounds for the generative approach (top) and number of available and high-scoring compounds in the in-house library (bottom) for each exercise. Percentages represent the proportion of total compounds meeting the high-score thresholds. **(c)** Similarity analysis of high-scoring compounds within each approach, illustrating the diversity of the candidate molecules for the generative (top) and library- based (bottom) approaches. **(d)** Similarity of high-scoring compounds to the active molecules in the training dataset, for the generative (top) and library-based (bottom) approaches.

Fig. 3b shows the number of unique molecules generated in each exercise, and the fraction with high scores for the dual-objective task, defined as those molecules with a predicted score above 0.6 and below 0.2 for the targeted and non-targeted cell line, respectively. For HEK293, we set the targeting at 0.8 instead, reflecting experimental observations that lower scores were less reliable in this cell line (Extended Data Fig. 6). Overall, the agent demonstrates low efficiency in generating high-scoring molecules (0.42%–3.29% of the total), but it still produced between 37 and 137 high-scoring candidates per exercise for further evaluation. Moreover, although the high-scoring generated molecules are quite chemically similar between them, with an average Tanimoto similarity around 0.85, they are also structurally distant from the closest active compound in the training set (average Tanimoto similarity around 0.3), showing the innovation potential of the generative models (Fig. 3c and d, top).

We also sought to investigate whether generative models show an added value over a more classical activity prediction strategy. Thus, we applied our prediction models to the IRB in-house library, which by this moment had been extended to 135,000 molecules, and followed the same validation procedure. In general, the number of high-scoring molecules found in each exercise was between 10 and 1,000 times lower than the ones generated by REINVENT (Fig. 3b, bottom). Similar to the generated molecules, the high-scoring compounds identified in our library showed a low chemical diversity, with slightly higher Tanimoto similarities around 0.9. However, as it could be expected, in this case many of these molecules were also close to the active compounds in the training (Fig. 3c and d, bottom).

Next, we conducted experimental validation of the generated and predicted high-scoring molecules. Since we could not afford the costs of synthetizing dozens of AI-designed new molecules, we searched for similar compounds to the high-scoring generated molecules in public catalogs, including ZINC20^44^, Mcule^45^, and ENAMINE REAL^46^ databases. For ZINC20 and ENAMINE, we used the SmallWorld Search^47^ to identify the closest matches based on graph edit distance (GED). Additionally, we employed the ENAMINE and Mcule search engines to find structurally similar molecules using Tanimoto similarity. For each query we retained up to 10 closest matches with a Tanimoto similarity >0.7 or GED <5, and evaluated these compounds using our prediction models. We identified at least one similar analogue to the generated molecules for all the seven exercises, but in four of them we could only find one suitable compound that passed all the stringent high-scoring criteria (Fig. 4a). Supplementary Table 9 shows the number of high-scoring molecules for each exercise, the number of close analogues found, and the ones that met the high-scoring criteria can be found in the GitHub repository (see *Data Availability*). Finally, we purchased the qualifying molecules, between 1 to 22 per exercise, and tested them experimentally. Further details about the search and selection processes are provided in *Methods*. Additionally, we tested all the high-scoring compounds identified in the in-house library, ranging from 2 to 27 compounds depending on the exercise (Fig. 4d; Supplementary Table 10).

**Fig. 4.**
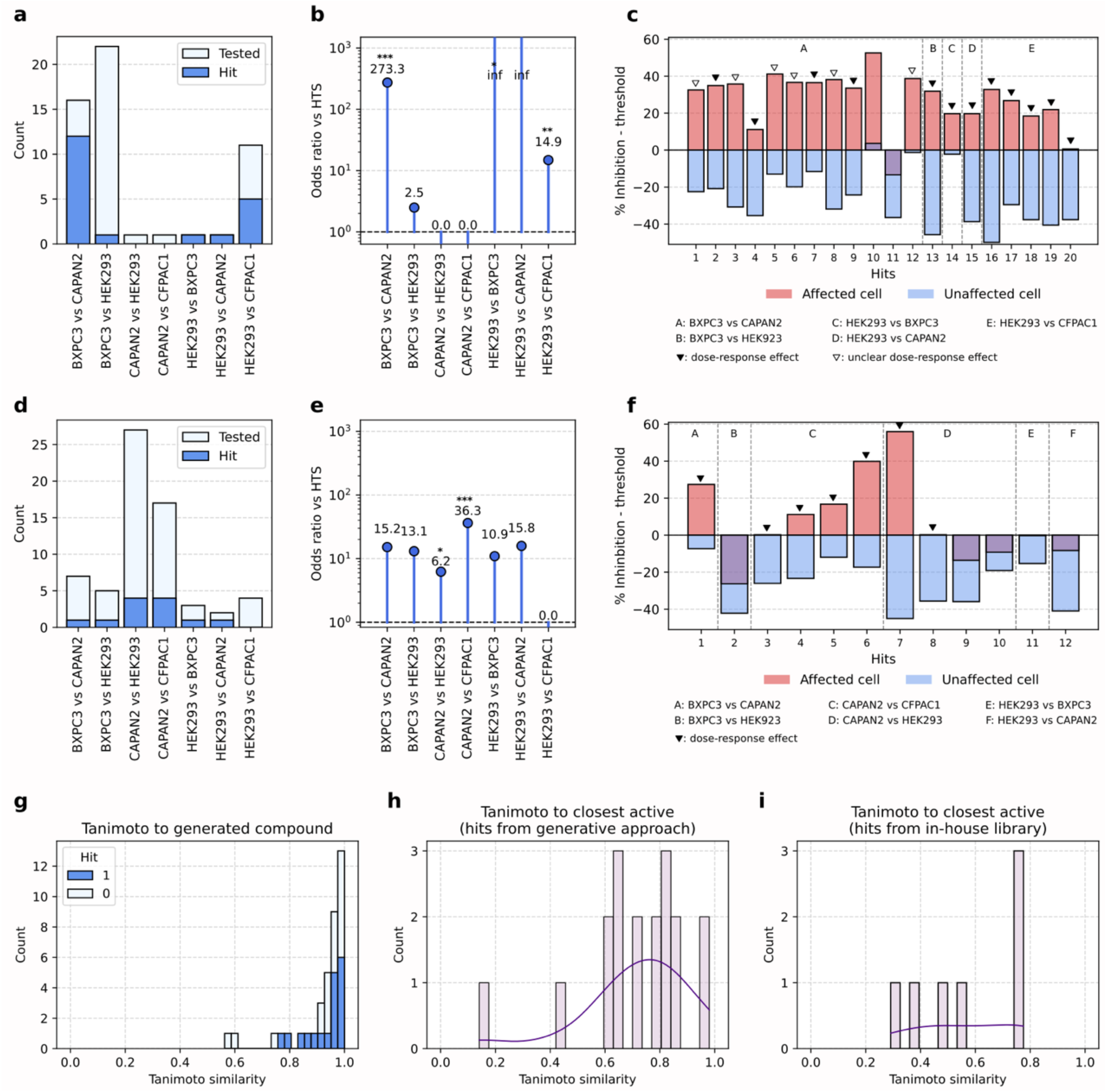
Experimental evaluation of generation and prediction-only approaches. Panels a-c correspond to the generative approach, while panels d-f correspond to prediction-only approach. **(a)** Number of generated compounds tested and the subset that showed the desired selective cytotoxic effects (hits) for each exercise. **(b)** Enrichment of hits identified through the generative approach compared to the HTS (quasi-random selection). *, **, ***: p-value < 0.05, 0.01, 0.001. **(c)** We performed the confirmation of hits identified in (a) through a dose-response (DR) assay. The plot shows the % inhibition at 10 μM for hit molecules in the generative approach for each exercise, relative to the cytotoxicity thresholds for each cell line. This representation highlights how much effect the compounds exhibit above or below the thresholds. Triangles indicate compounds for which a dose-response relationship was observed in the target cell lines. DR curves are available in Extended Data Fig. 8. **(d)** Number of high-scoring compounds from the in-house library tested (predictions-only approach) and the subset that showed the desired selective cytotoxic effects (hits) for each exercise. **(e)** Enrichment of hits identified through the prediction-only approach compared to the HTS (quasi-random selection). **(f)** Same as (c) but for hits identified through the prediction-only approach. DR curves are available in Extended Data Fig. 9. **(g)** For compounds tested in the generative approach, Tanimoto similarity between the generated compound and the final compound tested, and whether the compound was active (1) or not (0). **(h)** Tanimoto similarity of hits identified through the generative approach to the closest active compound in the training set. **(i)** Tanimoto similarity of hits identified through the prediction-only approach to the closest active compound in the training set.

Fig. 4 presents the results of the experimental validation, where we tested the compounds in triplicate at 10 μM, followed by a dose-response curve of those molecules that matched the desired effects. We tested 45 analogues to the AI-designed molecules, and 20 of them showed the desired cell-specific cytotoxic effects. Indeed, we validated at least one compound in 5 of the 7 exercises tested, with 4 of them showing a very significant increase in hit discovery rate compared to the HTS (odds ratio 14,9 - οο, p-value < 0.01; Fig. 4b). The hit compounds were structurally diverse and showed a wide range of chemical similarity with the active compounds in the training set (Fig. 4h). We further validated the hit compounds in dose-response assays (Fig. 5a, Extended Data Fig. 8), confirming 18 out of the 20 molecules tested. Among the confirmed hits, we successfully calculated IC50 values (at least in the targeted cell line) for 12 of the 18 molecules, ranging 0.97 to 16.6 μM. Next, we assessed whether the distance between the purchased and tested molecules to the original design was related to the likelihood of exhibiting the desired effect on the pair of cell lines. We observed that molecules more structurally distant from the generated candidates were less likely to exhibit the desired activity. Hits tended to be closer to the generated molecules, indicating that structural fidelity to the generated design is important for maintaining the predicted activity (Fig. 4g). These findings suggest that our results could improve further if we were able to synthesize and test the exact molecules produced by the generative model instead of the closest analogue in public vendors. We also tested 56 small molecules from the 135,000 compounds in the IRB library that scored high in the predictive models for each exercise using the same procedure. In this case, we identified a total of 12 hits showing the expected cell specificity, validating at least one compound in 6 of the 7 exercises. We also observed a significant increase in hit discovery rate with respect to HTS in 2 cases (odds ratio 6,2 - 36,3, p-value < 0.05; Fig. 4e). These hit compounds also showed chemical diversity with the active compounds in the training set (Fig. 4i). In the dose-response assays we confirmed 7 of the 12 hits, with IC50 values ranging from 5.9 to 15.1 μM (Fig. 4f, 5b and Extended Data Fig. 9).

**Fig. 5.**
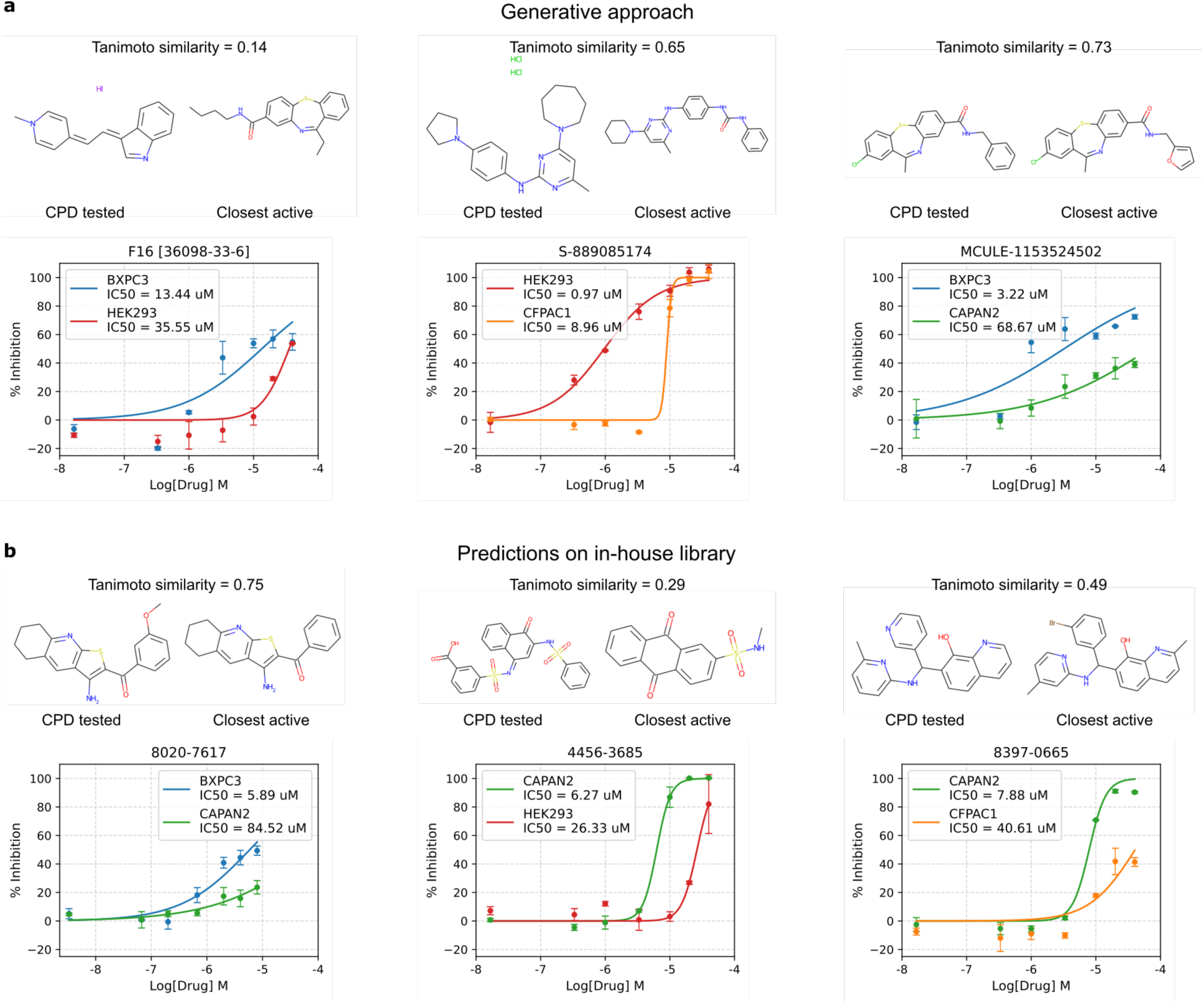
Dose-response curves for exemplary hits in the generative and prediction-only approaches. The hits from both approaches were further validated through a dose-response assay. **(a)** Three exemplary hits for the generative approach with good dose-response effects and different levels of similarity to the closest active in the training set, coming from different exercises. **(b)** Same as (a) for the prediction-only approach.

Overall, when comparing the two strategies, we observed that the two approaches are able to identify a set of diverse molecules with the desired cell specificity, finding at least one hit for most exercises, with a success rate significantly higher than blind HTS. However, the fraction of hits over the tested molecules, and more importantly the dose-response validation, is markedly higher in the AI-designed molecules. Moreover, since the baseline used for comparison –the HTS–already contains more hits than what would be expected in a blind assay (Supplementary Table 4), the calculated hit enrichments likely underestimate the relative improvement achieved with both the generative and the predictive strategies.

## Discussion

In this study, we explored whether generative artificial intelligence could be harnessed to design small molecules with cell-selective cytotoxic activity, moving beyond the conventional paradigm of target-based discovery. Our approach combined three complementary pillars: first, the generation of a comprehensive in-house cytotoxicity dataset by screening over 11,000 molecules across a diverse panel of pancreatic cancer lines of therapeutic relevance^48–50^ and control cells; second, the development of robust machine learning models trained on small-molecule bioactivity descriptors, capable of predicting cytotoxic responses with strong performance; and third, the integration of these predictive models into a reinforcement learning–based generative framework (REINVENT) to design novel chemical structures optimized for desired selectivity profiles. Our results demonstrated the feasibility of using genAI for phenotype-based drug design, and our strategy also resulted in the identification and experimental validation of novel molecules exhibiting the predicted selective cytotoxicities.

Our findings highlight several remarkable advances. By creating a high-quality cytotoxicity dataset across six pancreatic cancer cell lines with distinct mutational landscapes, complemented by two auxiliary lines representing non-tumorigenic and unrelated tumor contexts, we established a reliable foundation for training predictive models. Importantly, we showed that models based on bioactivity signatures consistently outperformed traditional fingerprint-based representations, highlighting the added value of incorporating systems-level chemical and biological information. Moreover, the predictive models proved effective not only retrospectively but also prospectively, enriching hit discovery in unseen compounds from our extended library. More critically, by embedding these predictors into a generative framework, we were able to design molecules tailored to cell-specific profiles, including challenging cases such as distinguishing between two pancreatic cancer lines with highly similar transcriptomic and mutational signatures. Experimental validation confirmed the robustness of the approach: AI-generated molecules exhibited consistently higher hit rates and selectivity compared to random screening, demonstrating the translational potential of coupling predictive and generative models.

Despite the promising outcomes, our study also highlighted several limitations. First, while the generative framework succeeded in producing compounds with the desired predicted profiles, its efficiency was relatively low, with only a small fraction of generated molecules meeting the dual-objective criteria. There is thus room for methodological improvement, for example, by sampling from intermediate training checkpoints, employing curriculum learning strategies more systematically, or integrating uncertainty-aware reward functions. Second, experimental validation was limited by practical constraints in accessing or synthesizing the exact AI-generated molecules. Instead, we relied on close analogues sourced from commercial vendors, which, although successful in multiple cases, likely underestimated the true potential of the generative designs. Direct synthesis of AI-proposed compounds, or restricting generative search spaces to synthetically accessible ultra-large libraries such as Enamine REAL^46^, could address this gap in future work. Additionally, while we confirmed selective cytotoxicity in multiple exercises, our experimental validation was limited to short-term cell viability readouts at micromolar concentrations, as these were the conditions in which the training data was generated. Further work could extend this validation to more detailed pharmacological characterizations, including response effects. Note that our work remains primarily proof-of-concept, demonstrating the feasibility of designing selective phenotypic modulators, but we never intended to achieve fully optimized drug candidates.

Looking forward, one immediate direction is to enhance the generative process by integrating multi-omics data directly into the optimization loop, leveraging transcriptomic or proteomic bioactivity signatures available in the Chemical Checker^27,29^. This would help generative models to navigate cellular states rather than relying solely on pre-trained cytotoxicity predictors. Another exciting possibility is to couple generative chemistry with causal inference frameworks to disentangle the drivers of selective cytotoxicity, thereby improving interpretability and guiding more rational optimization. Finally, expanding our approach from single-task selectivity to multi-objective optimization, including pharmacokinetics or toxicity predictors, would bring the strategy closer to real-world drug development needs.

Overall, our study demonstrates that deep generative models can be used not only to optimize physicochemical properties or protein-binding affinities, as in most previous applications, but also to design molecules with complex phenotypic outcomes such as cell-specific cytotoxicity. By integrating high-quality screening data, machine learning predictors, and reinforcement learning– based molecule generators, we successfully identified new compounds with selective activity against pancreatic cancer cell lines, validated experimentally. While challenges remain in terms of efficiency, synthetic accessibility, and translational validation, this work establishes a blueprint for a new generation of phenotype-driven drug discovery approaches that are target-agnostic yet biologically informed. As methods mature and experimental pipelines become more tightly integrated with generative design, we anticipate that this synergy between AI and phenotypic screening will open novel avenues for discovering first-in-class therapeutics against diseases with high unmet clinical need, such as pancreatic cancer.

## Methods

### In-house library of compounds

The original IRB library (in-house library) comprised 151 plates containing 47,489 compounds (∼320 per plate), sourced from ChemDiv (San Diego, CA) diversity libraries, and it was stored in 384-well plates at −80°C. During the course of the project, the in-house library was extended to include a total of 144,684 molecules, following the same logic of maximising chemical diversity. The additional compounds came from the same catalogue. This later version is referred to as ‘extended in-house library’ in the text.

### Compound descriptors (ECFP4 and bioactivity Signaturizers)

We computed classical ECFP4 descriptors using rdkit version 2023.03.2, with radius=2, bit size=2048. We obtained the CC bioactivity descriptors using the version 1.7 of the Signaturizers software^28^. In brief, the Signaturizers are a collection of deep learning models that predict the bioactivity signature of any given compound in 25 different bioactivity spaces defined in the ChemicalChecker^27^, from physicochemical properties to clinical outcomes, including also target binding or gene expression effects. Each of the different 25 Signaturizer descriptors have a length of 128, and the GLOBAL descriptor concatenates all of them, with a final dimension of 3,200 (CC_GLOBAL in the text).

### Plate selection for HTS

To maximise number of potential hits in the high-throughput screening (HTS), we used cell line sensitivity data from DrugCell^39^, which integrates information from the Cancer Therapeutics Response Portal (CTRP) v2^37^ and the Genomics of Drug Sensitivity in Cancer (GDSC)^38^ database. We analysed eight cell lines: the six pancreatic cancer cell lines studied in this paper (BXPC3, CAPAN2, CFPAC1, HPAFII, PANC10.05, and SW1990), the AsPC-1 cell line (initially part of the study but later excluded due to cell identity issues), and one osteosarcoma control (U2OS). We focused on 398 compounds tested on all the 8 cell lines.

#### Binarization of Sensitivity Data

We binarized the drug sensitivity data from DrugCell (AUC) using the waterfall method first described by Barretina et al.^51^, and used since then in different subsequent works (e.g.,^52,53^). This method ranks cell lines by response and defines a sensitivity threshold based on the plot shape. We applied the method to each compound, keeping 1-20% of sensitive lines with a maximum AUC of 0.9.

#### Neighbour identification

We defined five groups of compounds based on the sensitivity profiles retrieved from DrugCell:

● Active in 1 cell line (n = 47)
● Active in 2 cell lines (n = 31)
● Active in all cell lines (n = 6)
● Active in all pancreatic cancer lines but not in U2OS (n = 4)

We identified neighbours for these 100 compounds within the IRB library using three Signaturizer descriptors (see *Methods, Compound descriptors*): A4 (MACCS keys), B4 (binding), and D2 (cell line sensitivity). We identified neighbour compounds based on cosine similarity, leveraging the CC pre-established threshold corresponding to a p-value, calibrated against the distance distribution of the 1 million molecules within the CC resource for each descriptor. This yielded 60, 216 and 3,919 neighbours based on p-value < 10e-5, 10e-4 and 10e-3 respectively (closest, medium and distant neighbours).

#### Plate selection

We ranked the plates by the number of neighbours they contained, weighted by proximity (closest = 100, medium = 10, distant = 1). We selected the first 35 plates, encompassing 11,053 molecules (100% of closest neighbours, 38.4% of medium and 28.4% of distant ones). Molecules in the selected plates originated from all criteria and descriptors.

### Cell culture, treatments and cytotoxicity assays

We acquired the pancreatic cancer cell lines BXPC3, CAPAN2, HPAFII, PANC10.05, and SW1990 from the American Type Culture Collection (ATCC), and CFPAC1 and AsPC-1 lines were kindly provided by Dr Cristina Mayor-Ruiz (IRB Barcelona). We maintained CAPAN2 cells in McCoy’s 5A medium, PANC10.05 in RPMI medium supplemented with 4 µg/ml insulin, CFPAC1 in IMDM, HPAFII in EMEM, and AsPC-1, BXPC3 and SW1990 in RPMI medium. All media were supplemented with 10% fetal bovine serum (FBS), 100 U/ml penicillin, and 100 µg/ml streptomycin. Additionally, U2OS and HEK293 cell lines were kindly provided by Dr Jens Lüders (IRB Barcelona), and cultured in DMEM supplemented with 10% FBS, 100 U/ml penicillin, and 100 µg/ml streptomycin. All cells were grown at 37°C in 5% CO₂ and 90% humidity.

We first tried to reproduce the drug sensitivity data from DrugCell on 12 previously reported, selective drugs against our cell line panel. We performed dose-response MTT experiments measuring inhibition in cell viability. We seeded 100 µl of cells in 96-well clear flat-bottomed plates at the following densities: BxPC-3, CFPAC-1, HPAF-II, SW1990, HEK293 and U2OS 10.000 cells/well; Capan-2 and Panc 10.05 12.000 cells/well and we kept them at 37°C for 24h before adding drugs. The following day, we added 20 µl of compound (6x) to each well to create an 8- point concentration-response curve. 48h after compound addition, we added 0.5 mg/ml MTT solution to each well and we incubated cells for 3h at 37°C. Then, we removed supernatant with a multichannel pipette and dissolved formazan salts with 140 µl DMSO. We incubated the plates for 10 minutes in a shaker at RT and read at 570 nm using an ELx808 plate reader (Agilent Biotek). We measured % inhibition in viability by normalizing each compound with the positive control compound (Sepantronium bromide, a survivin inhibitor that strongly reduces tumor cell viability and induces apoptosis). We analyzed dose-response curves using Graphpad Prism 9 (San Diego, CA, USA).

To perform the HTS, we conducted a viability-based assay, with cell concentrations optimised to maintain logarithmic growth throughout the 3-day experiment. We seeded the cells in 384-well white plates at the following densities: BXPC3, CAPAN2, and PANC10.05 at 1000 cells/well; HPAFII and SW1990 at 750 cells/well; CFPAC1 at 500 cells/well; and AsPC-1 at 350 cells/well. After 24 hours at 37°C, we added the ∼11,000 compounds to the cells, distributed across 35 plates, using an Echo 650 dispenser (Beckman Labcyte) to achieve a final concentration of 10 µM in a single replicate. We dissolved the compounds not belonging to the IRB library in DMSO to a concentration of 10 mM, aliquoted, and stored at −80°C until use. We strategically positioned positive (Sepantronium bromide) and negative (vehicle) controls on each plate, and we maintained the DMSO concentration below 1% to avoid adverse effects on cell growth.

After 48 hours, we added 30 µl of CellTiter-Glo solution (Promega) directly to 30 µl of cells to each well (1:1 ratio). The CellTiter-Glo® detection kit assesses cell viability by measuring ATP levels through a luciferase-based luminescent reaction. We measure luminescence using an Envision plate reader (Revvity) following a 10-minute incubation at room temperature. We calculated the percentage inhibition in viability by normalizing the luminescence signal to vehicle-treated wells (100%) and sepantronium-treated wells (0%). A Z-prime score above 0.5 ensured quality control for individual plates. For confirmation, we retested the ∼750 active hits identified in the HTS in the same seven pancreatic cancer cell lines and two additional non-pancreatic cell lines (HEK293 and U2OS). We seeded the HEK293 and U2OS cells at 750 and 1000 cells/well, respectively, with the same densities as before for pancreatic cell lines. We performed confirmation experiments at 10 µM in duplicate, and we compared the results to the initial HTS for data replication. After excluding AsPC-1 because of cell identity issues, we confirmed 392 hits from the 751, demonstrating consistent activity across experiments. For the final dataset, we considered the 3 available replicates (1 from the HTS and 2 from the confirmation experiment) and computed the mean of the closest 2 as the final % inhibition of each compound in each cell line. In the case of HEK293 and U2OS, we simply considered the mean of the 2 available replicates. All the subsequent cytotoxicity experiments to validate predictions or generated compounds were performed following the same protocol. We used the results of the HTS as random expectation for hit identification in all the PC cell lines and for the HEK293 and U2OS cells, since they were not included in the HTS, we re-tested one plate of the HTS to obtain a baseline.

Additionally, we created dose-response curves to evaluate and compare the effects of the compounds identified by our algorithm with those predicted from the IRB library. To achieve this, we exposed the cells to a series of concentrations spanning a broad dynamic range for 48 hours, following the previously described experimental protocol. We normalized the dose-response curves to the control curves obtained with sepantronium. We applied a four-parameter logistic (variable-slope) inhibition model for curve fitting using the scipy.optimize.curve_fit function in Python. This experimental setup allowed for the determination or prediction of the half-maximal inhibitory concentration (IC50) for each compound, providing a quantitative measure of their potency and enabling a direct comparison between algorithm-identified compounds and IRB library predictions.

#### RNA-seq analysis

We acquired QIAGEN RNA extraction kits and followed the specified RNA extraction protocol, obtaining the RNA from the seven pancreatic cancer and auxiliary cells. We conducted RNA sequencing using a poly-A selection protocol, employing a paired-end 150 bp format with 25 million reads per sample on a Novaseq S4 platform. First we trimmed the reads to remove adapter sequences using Trim Galore^54^. We aligned cleaned reads to the SILVA rRNA database^55^ using Bowtie2 (v2.2.2)^56^ with the options -q -t -N 1 -k 1 -p 10 to remove rRNA-derived reads. We subsequently aligned rRNA-unaligned reads to the hg38 genome using STAR^57^ (v2.7.10a) with the parameters --outFilterMismatchNoverLmax 0.05 --outFilterMatchNmin 25 and the rest left to default. We then computed counts per genomic feature with the R package Rsubread^58^ using the featureCounts function with strandSpecific=2 and countMultiMappingReads=TRUE.

### Correlation between cell line sensitivity and other cell line descriptors

We represented each cell by a sensitivity vector, capturing the percentage inhibition of the 751 compounds tested in the confirmation experiment. We repeated the same approach using other cell line descriptors: gene expression vectors derived from the RNA-seq data experiment, binary mutational descriptors from DrugCell^39^, proteomics data from DepMap^42^ and several 11 other descriptors from the Bioteque^25^ web resource (Supplementary Table 11). To assess similarity between cells, we employed Euclidean distance for continuous values or Jaccard’s similarity index for binary ones.

Finally, we computed the Spearman’s correlation coefficient between the distance found using the sensitivity data vs each of the other modalities in order to evaluate the concordance between cell similarity patterns in sensitivity and other cell line descriptors.

### Cell-line sensitivity prediction

#### Input data

We binarized the data from the confirmation experiment using thresholds derived from the HTS. Within each cell line, a compound was labelled "active" if its % inhibition was above the mean % inhibition plus 2.5 standard deviations of all compounds tested during the HTS for that cell line. The thresholds are therefore cell line-specific.

We then compared the performance of 2 types of representations for the molecules: ECFP4 fingerprints^43^ and the GLOBAL Signaturizer descriptors^28^ (see *Methods, Compound descriptors*). For these GLOBAL descriptors (CC_GLOBAL) we applied the k-best dimensionality reduction algorithm from scikit-learn to get back to the 128-length vectors from the original individual descriptors, which would help avoid overfitting and obtain lighter models.

#### Model training and evaluation

We trained cell line-specific ExtraTrees classifiers in a stratified 10-fold cross validation setting, using random splits and with a fixed positive:negative ratio of 1:10. For selecting the negative samples for each cell line, we randomly picked inactive compounds from the confirmation experiment and, whenever this was not enough to fulfil the 1:10 ratio, we picked extra inactive molecules randomly from the HTS. The HEK293 classifier directly used the available confirmation experiment data without enforcing the 1:10 ratio, due to the high number of active compounds in the confirmation experiment and the lack of data from the HTS.

We re-trained the final prediction models considering all the data from the different splits, using the CC_GLOBAL descriptors from the Signaturizers.

#### Selection of compounds for experimental validation

To experimentally validate the sensitivity predictors for the 6 pancreatic cancer cell lines, we used the classifiers to make predictions in the whole original IRB library, removing the ones used in the HTS and confirmation assays. We divided the remaining compounds in 4 groups, according to their Tanimoto similarity to the closest compound in training: <0.4, 0.4-0.6, 0.6-0.8, >0.8, and we chose the top 7 scoring compounds in each range for each cell line up to a total of 320 compounds (Extended Data Fig. 5). The HEK293 were also included in the experimental validation but not considered in the selection process.

### Generative models

For each gen AI cell-specific exercise, we first attempted to run REINVENT 3.2^10^ with the classical reinforcement learning (RL) configuration. The exact parameters used are specified in the Supplementary Table 8. In brief, we ran the models for 2,000 reinforcement cycles, with Scaffold Similarity as a diversity filter, and all the active compounds in training as inception molecules. In the scoring function, we included the property predictors for each of the cell lines with a sigmoid transformation of the output score to maximise high scores and penalise mediocre ones. We also included the default custom alerts and a molecular weight filter of < 550Da as part of the scoring function.

In most exercises the classical reinforcement learning approach was not able to generate good-scoring molecules, we thus attempted a curriculum learning strategy. We started the curriculum phase optimising for molecules predicted to affect one of the cell lines, using the same parameters as in RL, but excluding the classifier for the second cell line from the scoring function. Then, once the algorithm reached a mean score of 0.6 for the generated molecules, we started the production phase, where we included the second classifier to the scoring function. Each phase ran for a maximum of 2,000 steps. Finally, in some exercises we observed a repetitive substructure (Extended Data Fig. 7) appearing in most of the generated molecules. We decided in these cases to run the curriculum learning models with the same parameters, but adding to the scoring function a filter to penalize structures containing this motif.

After running all the models, we kept those that reached a mean score of at least 0.4, and we sampled 10,000 molecules per exercise for further evaluation. We defined high-scoring molecules as those with a predicted score above 0.6 for the targeted cell line and below 0.2 for the non-targeted one. For HEK293, the threshold for targeting was set at 0.8 instead, reflecting experimental observations that lower scores were less reliable in this cell line.

### Search for generated compounds in public libraries

For all the high-scoring generated compounds in the different exercises, we looked for either the exact molecule or a close analogue in different public libraries. More specifically, we used the search engine SmallWorld Search^47^ to quest ZINC20 for sale subset (2022 Q1, 1.6B) and REAL Database (ENAMINE, 2022 Q1, 4.5B). We searched these databases based on the graph edit distance (GED). We selected molecules with a GED <= 5. Additionally, we also used the Mcule search engine^45^ to look for molecules in their catalogue. In this case the search was performed using Tanimoto similarity scores, and we selected molecules with a Tanimoto similarity ≥ 0.7. For all the identified compounds meeting our criteria, we ran again the cell-specific cytotoxicity predictors, and discarded the ones that did not pass the prediction thresholds defined in the previous section (*Methods, Generative models*). For each query molecule, we bought the closest, high-scoring compound found, if any (Supplementary Table 10).

## Data availability

All the experimental data corresponding to compound screenings (HTS, validation of cytotoxicity predictions and validation of generated molecules) and RNA-sequencing are available in the Supplementary Tables and at https://github.com/sbnb-irb/cytotoxics_design.git.

## Code availability

Cytotoxicity prediction models and code to run new predictions is available at https://github.com/sbnb-irb/cytotoxics_design.git.

## Acknowledgments

We would like to thank all public databases and methods that have allowed this study, and the members of the Structural Bioinformatics and Network Biology lab from IRB Barcelona for the constant discussions.

## Funding

PA acknowledges the support of the Generalitat de Catalunya (2021 SGR 00876), the Spanish Ministerio de Ciencia, Innovación y Universidades (PID2023-152296OB-I00), and the European Commission (CLARITY: 101137201). GR-G is a recipient of an FPI-SO fellowship (PRE2020- 092083. We also acknowledge institutional funding from the Spanish Ministry of Science and Innovation through the Centres of Excellence Severo Ochoa Award, and from the CERCA Programme / Generalitat de Catalunya.

## Author information

### Authors and Affiliations

Institute for Research in Biomedicine (IRB Barcelona), The Barcelona Institute of Science and Technology (BIST), Barcelona, Catalonia, Spain

### All the authors

Institució Catalana de Recerca i Estudis Avançats (ICREA), Barcelona, Catalonia, Spain Patrick Aloy

### Contributions

GR-G and PA conceived and designed the study, and wrote the manuscript. GR-G implemented all the computational models and analyzed the results. MS-S and JC performed the cell assays and dose-response studies. MB did the plate selection for the HTS. MC and IR ran the HTS in the IRB Barcelona Drug Screening core facility. All authors read and approved the final manuscript.

### Corresponding author

Correspondence should be addressed to Patrick Aloy.

## Ethics declarations

### Ethics approval and consent to participate

Not applicable

### Consent for publication

Not applicable

### Competing interests

The authors declare no competing interests.

## Extended data

**Extended Data Fig. 1.**
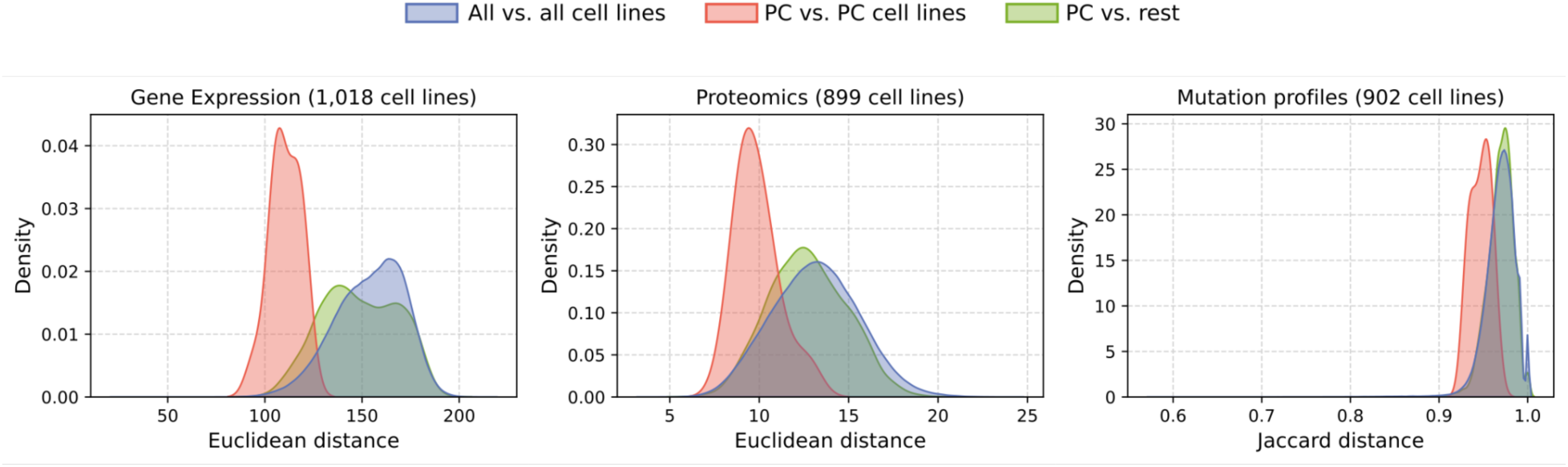
Distance between cell lines in different biological descriptors. *PC cell lines* correspond to the 6 pancreatic cancer cell lines included in this study. *All cell lines* are the total of cell lines for which the descriptors were available. Gene expression descriptors were taken from GDSC1000^59^ (RMA-processed basal expression levels for 17,737 genes measured in 1,018 cell lines). Proteomic descriptors were taken from DepMap^42^ (CCLE Reverse Phase Protein Array for 144 proteins in 899 cell lines). Mutational profiles come from DrugCell^39^, consisting of binary descriptors indicating mutated (1) or wild type (0) state for 3,008 genes in 902 cell lines.

**Extended Data Fig. 2.**
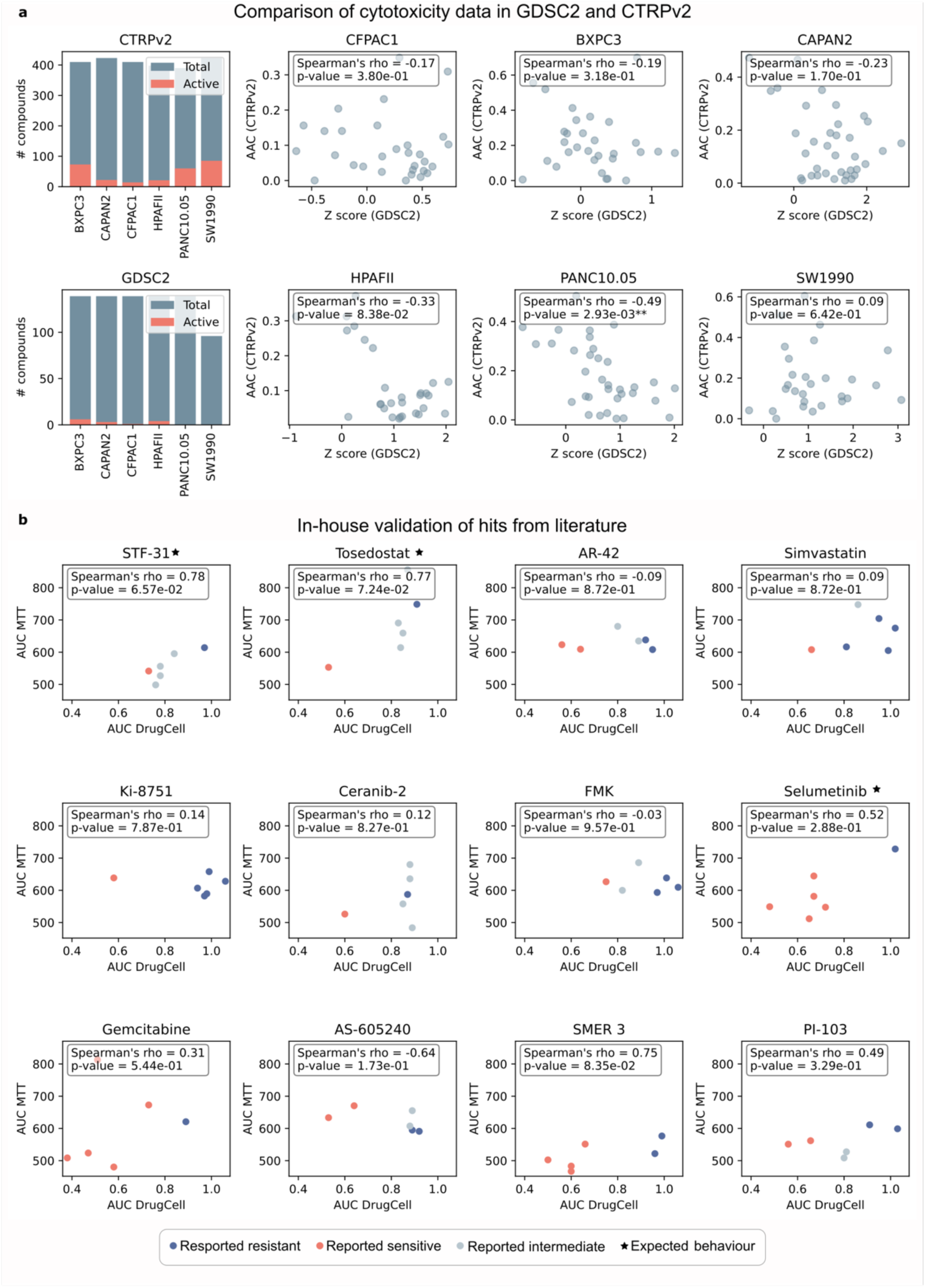
Comparison of data in different public databases and experimental confirmation for our cell line panel. **(a)** On the left, the number of compounds tested and those found to be active in CTRPv2^37^ (top) and GDSC2^38^ (bottom) for our pancreatic cancer cell lines. The scatter plots show the activity and correlation between overlapping compounds in both databases for each cell line. AAC: area above the dose-response curve. The larger the more sensitive. Z score: sensitivity z-score (computed per compound across cell lines). The smaller the more sensitive. Stars indicate significance levels (**: p-value < 0.01). **(b)** Experimental confirmation of selected molecules reported to have differential effects on our PC cell line panel. The scatter plots show the reported activity in DrugCell^39^ compared to the activity we observed. AUC: area under the dose-response curve. The smaller the more sensitive.

**Extended Data Fig. 3.**
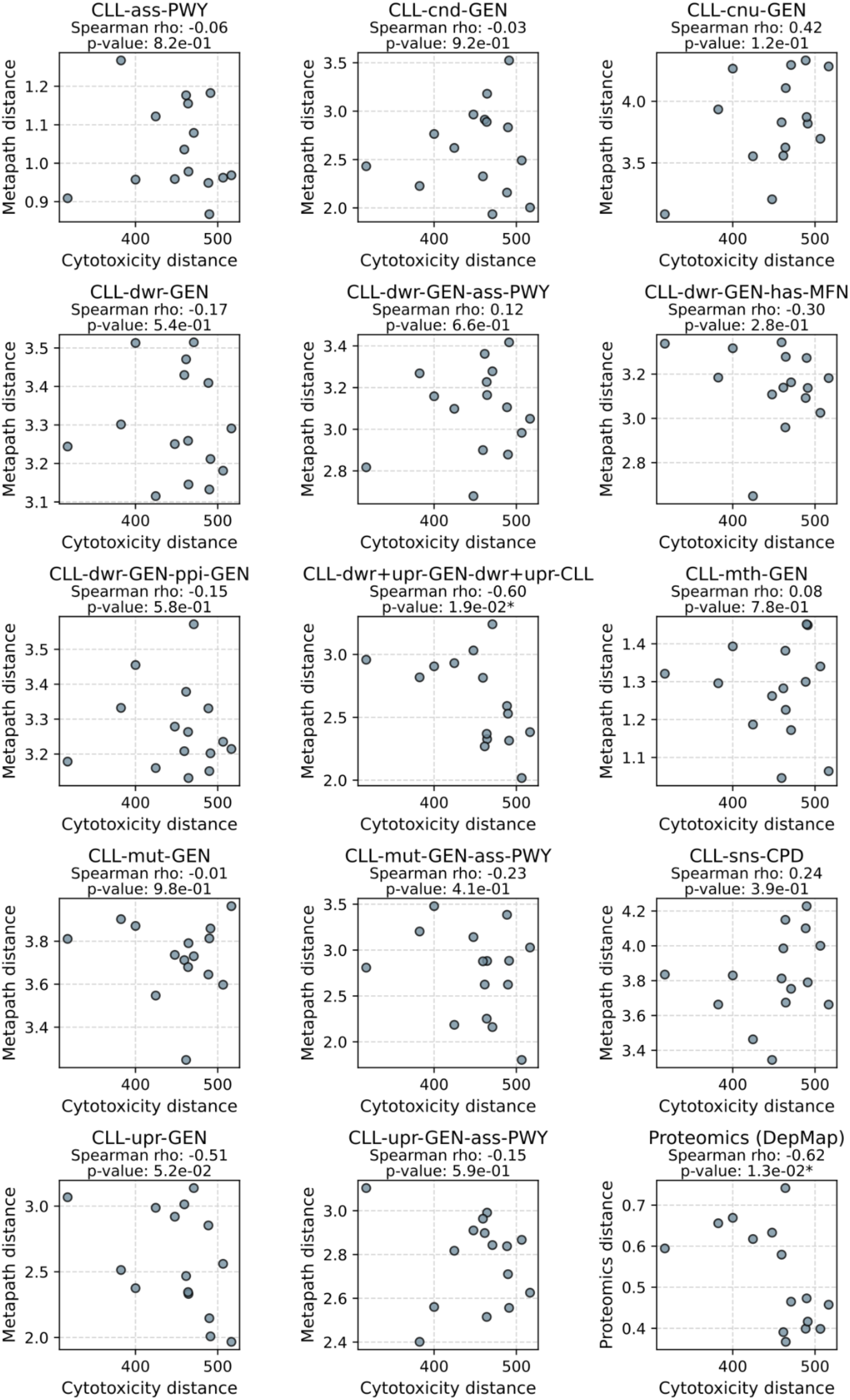
Comparison of cell line similarities in sensitivity vs. Bioteque metapaths and proteomics. Following the same principle as in Fig. 1e, we examined the correlation between cell line distances in cytotoxicity and distances in Bioteque metapaths^25^ and proteomics data from DepMap^42^, focusing on the six pancreatic cancer (PC) cell lines of interest (see *Methods*). Stars indicate the level of significance (*: p-value < 0.05). See Supplementary Table 11 for Bioteque metapath description.

**Extended Data Fig. 4.**
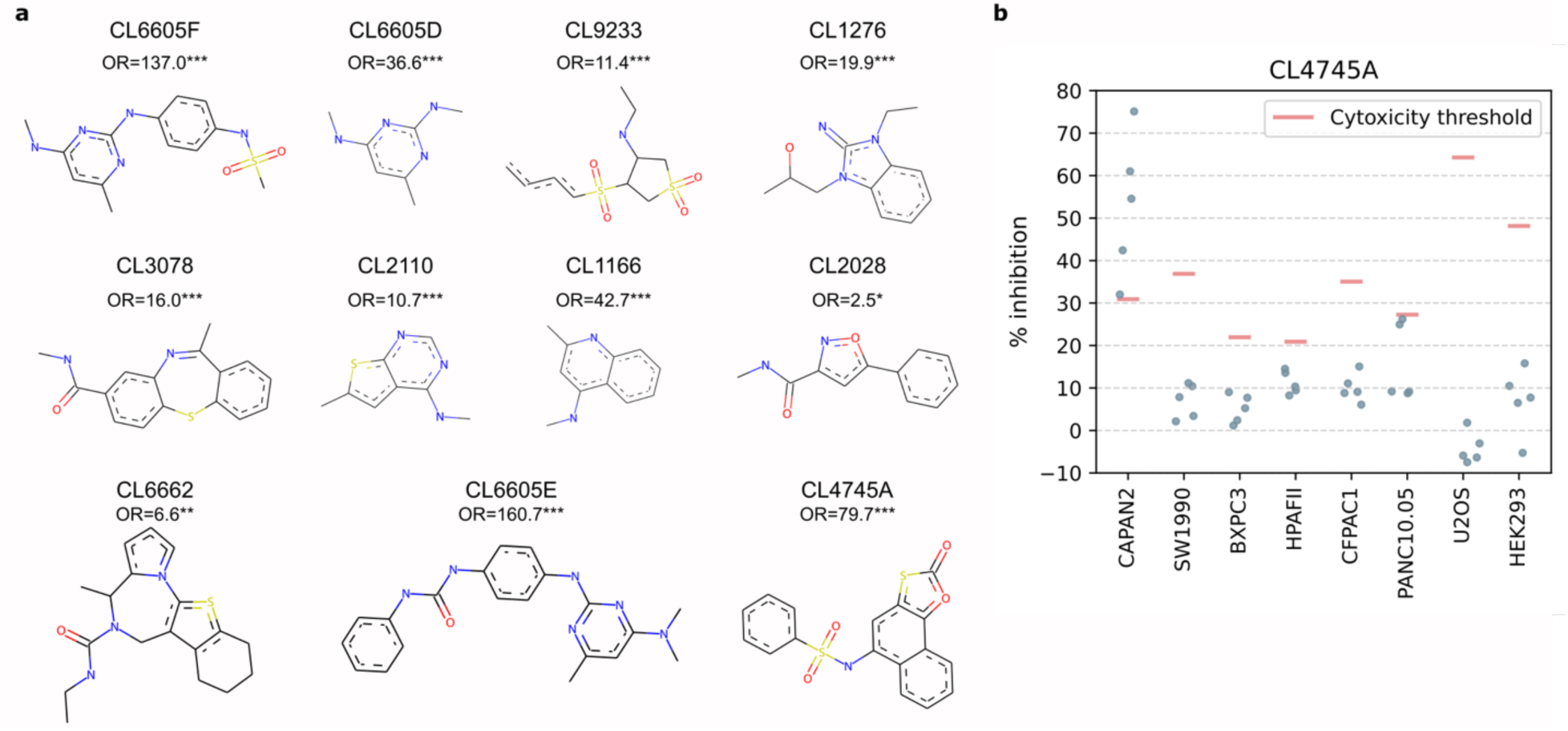
Scaffolds with >5 active molecules confirmed after HTS. **(a)** Molecular structures of the scaffolds with more than five active molecules confirmed in the high-throughput screening (HTS). OR: odds ratio from a Fisher’s exact test comparing ratio of active molecules containing those scaffolds compared to the whole HTS. *, **, ***: p-value<0.05, 0.01, 0.001. **(b)** % inhibition per cell line for the five molecules containing the scaffold CL4745A, which showed specificity to CAPAN2.

**Extended Data Fig. 5.**
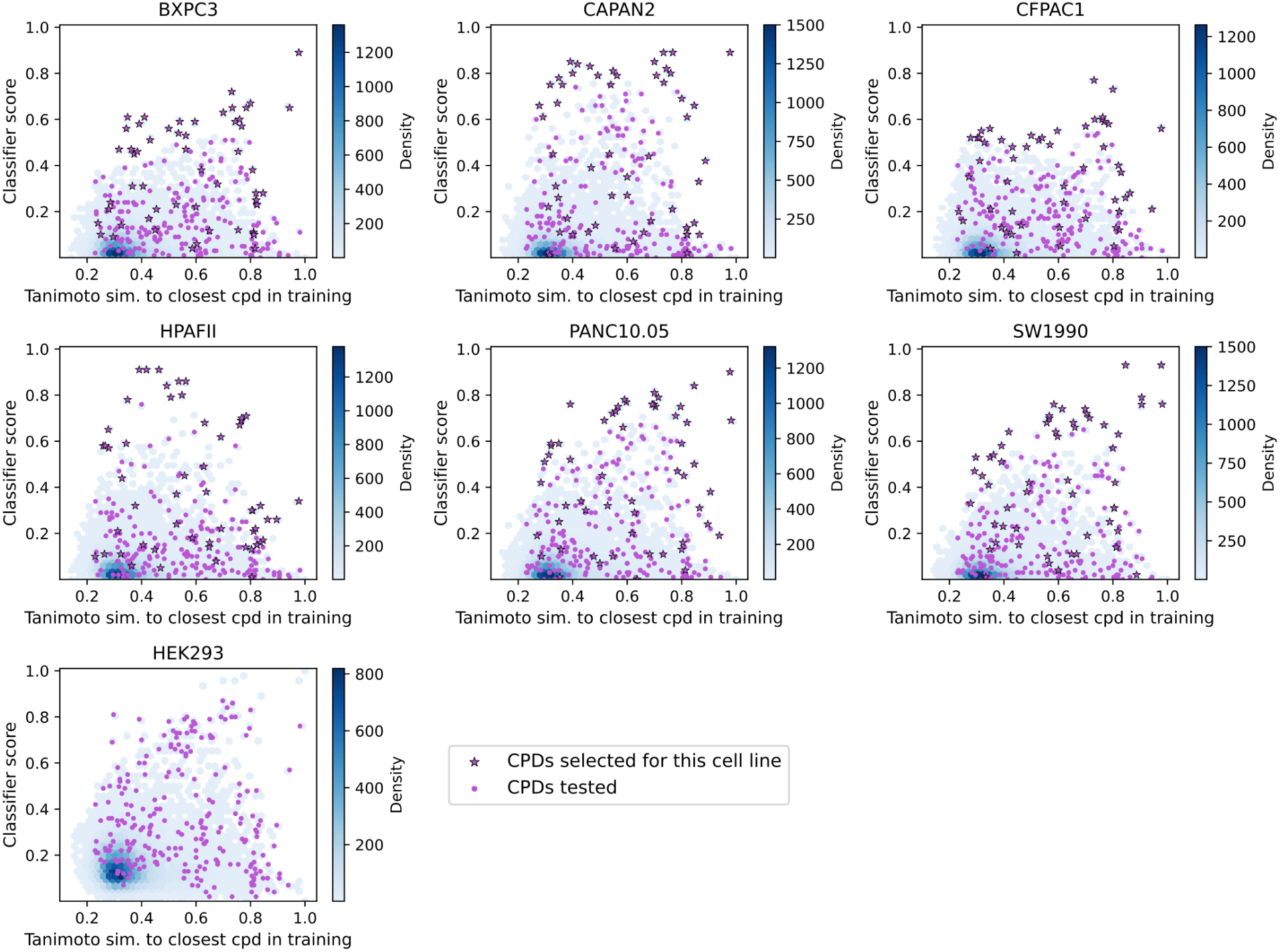
Tanimoto similarity to training compounds vs predicted score for compounds selected for validation of cytotoxicity predictors. To experimentally validate the cytotoxicity predictors, predictions were made for the entire in-house library, and a subset of compounds was selected based on varying Tanimoto similarity scores to the closest molecules in the training dataset and different classifier-predicted scores (see *Methods*). The plot shows, for each cell line, the Tanimoto similarities and predicted scores for the entire in-house library (blue) and highlights the selected compounds (pink). Star markers denote compounds specifically selected for the given cell line.

**Extended Data Fig. 6.**
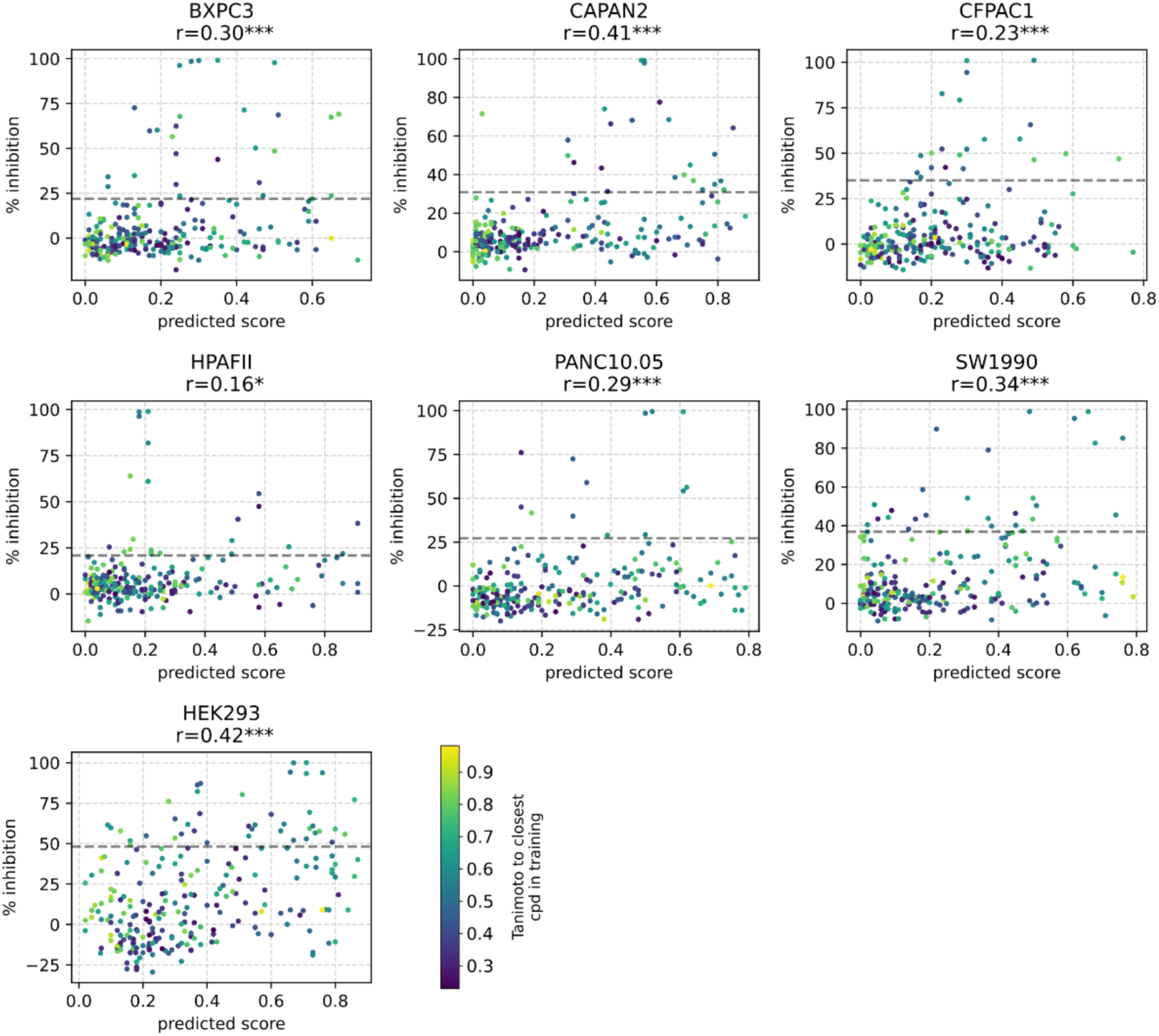
Experimental validation of the cytotoxicity predictors. Predicted scores versus % inhibition observed in the experimental validation of the cytotoxicity predictors for each cell line. Data points are colored based on Tanimoto similarity to the closest compound in the training dataset. Dashed lines represent the cytotoxicity thresholds. r shows Pearson correlation between axes. *, **, ***: p-value < 0.05, 0.01, 0.001.

**Extended Data Fig. 7.**
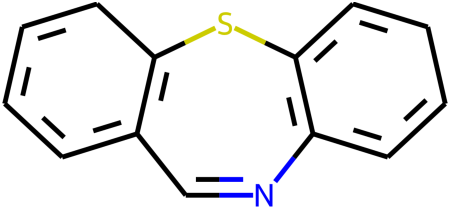
Repetitive substructure in the molecular generation process. Both for the exercises BXPC3 vs CAPAN2 and BXPC3 vs HEK293 we found this substructure to be extremely prevalent in the proposed designs, thus the generative cycle was run again penalising the molecules containing this motif.

**Extended Data Fig. 8.**
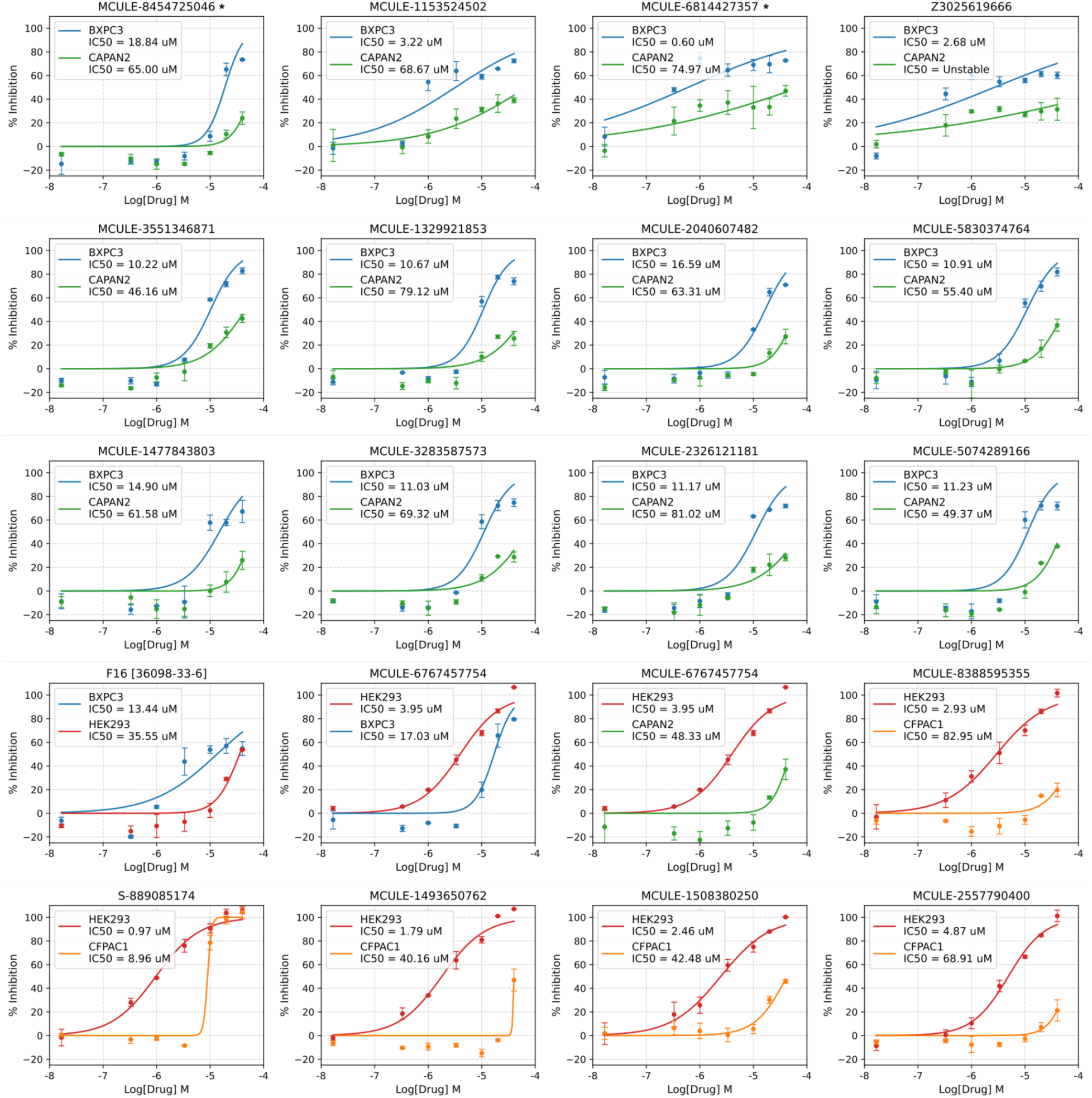
Dose-response curves for the hits found through the generative approach. Stars indicate compounds that did not pass the cytotoxicity thresholds at 10 μM in the dose-response assay.

**Extended Data Fig. 9.**
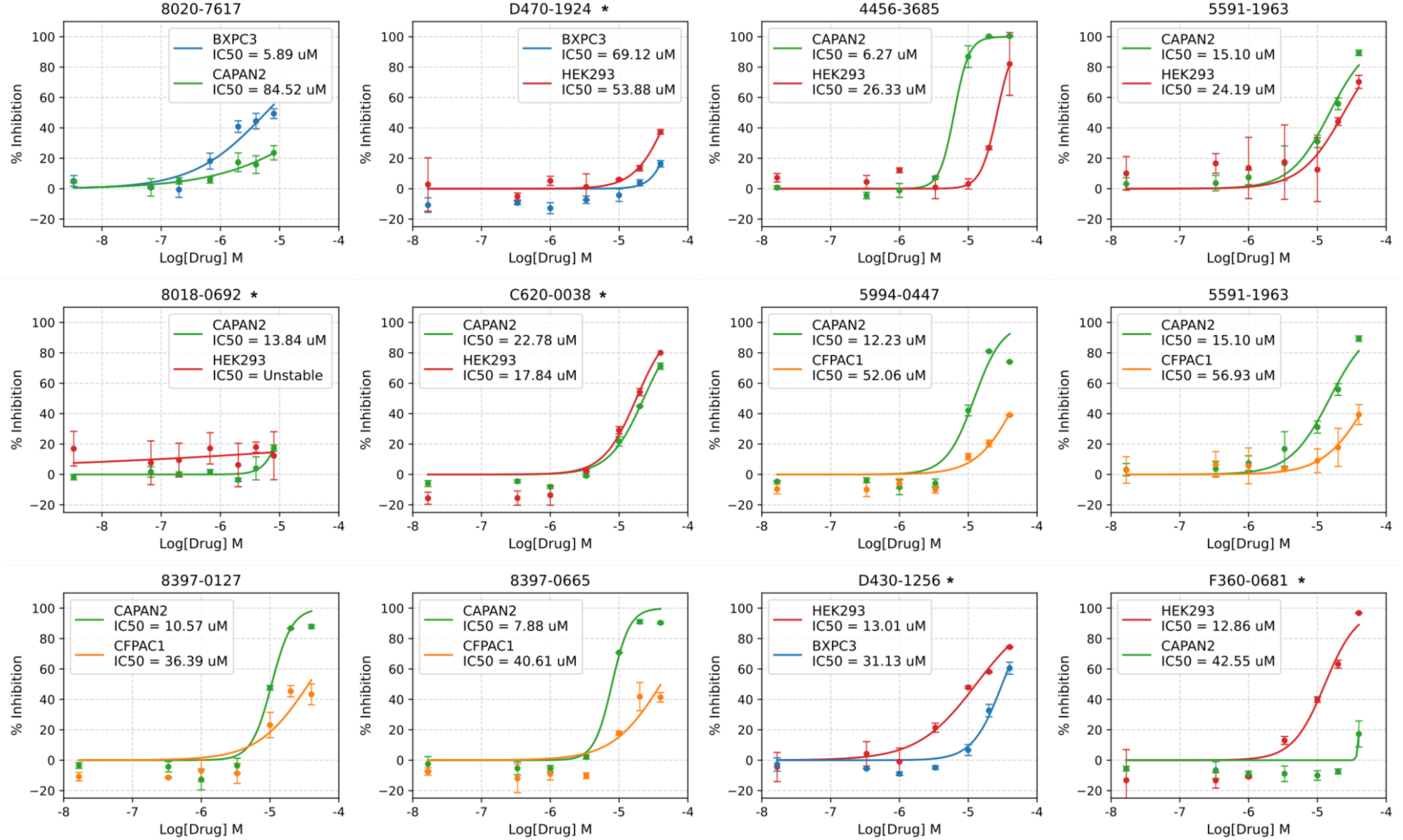
Dose-response curves for the hits found through the prediction-only approach. Stars indicate compounds that did not pass the cytotoxicity thresholds at 10 μM in the dose-response assay.

## Supplementary information

**Supplementary Table 1.**
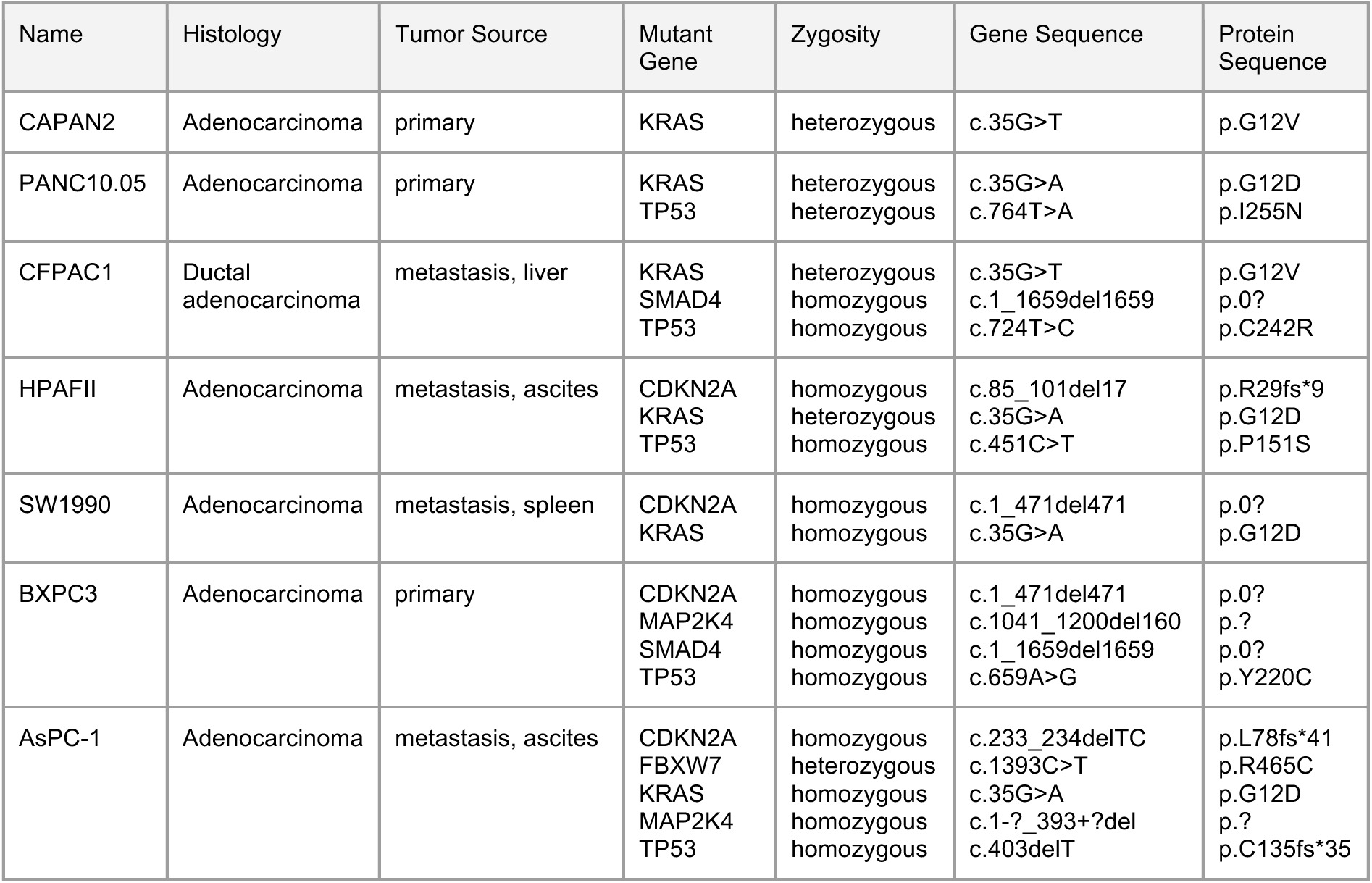
Description of pancreatic cancer cell lines (from ATCC® TCP-1026 Pancreatic Cancer Panel).

**Supplementary Table 2.** Cytotoxicity data in public databases.

| Database | # Compounds (CPD) | # Cell lines (CLL) | CPD-CLL pairs | # Pancreatic cell lines | # Cell lines of interest (9) | Info |
| --- | --- | --- | --- | --- | --- | --- |
| GDSC2 | 198 | 809 | 135,242 | 30 | 8 | IC50<br>IC50 Z-score<br>AUC<br>Sensitive/resistant classification |
| NCI-60 | 21,993 | 60 | 958,762 | 0 | 0 | Z-scored IC50 |
| CTRPv2 | 484 | 887 | 326,427 | 29 | 8 | AAC |
| DrugCell | 684 | 1,224 | 509,294 | 42 | 8 | AUC |

**Supplementary Table 3.** Evaluation of selective cytotoxicity in previously reported drugs. The table shows dose–response curves for 12 drugs with selective activity toward one or more cell lines in our panel, as reported in DrugCell.

**Supplementary Table 4.** Enrichment of hits within the nearest neighbors from reported actives in literature vs rest of molecules tested (HTS). ***: p-value < 0.001.

| Cell line | Odds ratio | p-value |
| --- | --- | --- |
| CAPAN2 | 2.26 | 8e-7*** |
| SW1990 | 2.37 | 4e-9*** |
| BXPC3 | 2.09 | 1e-5*** |
| HPAFII | 2.78 | 1e-9*** |
| CFPAC1 | 2.53 | 1e-9*** |
| PANC10.05 | 1.98 | 1e-6*** |

**Supplementary Table 5.** High-throughput cytotoxicity screening and hit confirmation. The table includes the final training set derived from this data, which we used to model cytotoxicity prediction, and the posterior screening of one of the plates from the HTS on HEK293.

**Supplementary Table 6.** Gene expression data. The table reports RNA-seq raw counts per cell line.

**Supplementary Table 7.** Experimental validation of cytotoxicity prediction models.

**Supplementary Table 8.** Parameters overview of REINVENT runs. *The aggregation method for the scoring function was *custom product* for all the exercises. **CLL1: targeted cell line. ***CLL2: spared cell line. The rest of the tunable parameters were kept as default.

| Exercises | Run type | Phase | # Epochs | Diversity filter | Scoring function* components (weight) |
| --- | --- | --- | --- | --- | --- |
| BXPC3 vs CAPAN2<br>CAPAN2 vs HEK293 | Reinforcement learning | - | 2,000 | Scaffold Similarity | - CLL1** classifier (5)<br>- CLL2*** classifier (2)<br>- Molecular weight (2)<br>- Default structural alerts (1) |
| BXPC3 vs CAPAN2<br>BXPC3 vs HEK293<br>CAPAN2 vs HEK293<br>CAPAN2 vs CFPAC1<br>HEK293 vs BXPC3<br>HEK293 vs CAPAN2<br>HEK293 vs CFPAC1 | Curriculum learning | Curriculum | Up to 3,000 | Scaffold Similarity | - CLL1 classifier (5)<br>- Molecular weight (2)<br>- Default structural alerts (1) |
|  |  | Production | 2,000 | Scaffold Similarity | - CLL1 classifier (5)<br>- CLL2 classifier (2)<br>- Molecular weight (2)<br>- Default structural alerts (1) |
| BXPC3 vs CAPAN2<br>BXPC3 vs HEK293 | Curriculum learning (unwanted moiety) | Curriculum | Up to 3,000 | Scaffold Similarity | - CLL1 classifier (5)<br>- Molecular weight (2)<br>- Default structural alerts (1) |
|  |  | Production | 2,000 | Scaffold Similarity | - CLL1 classifier (5)<br>- CLL2 classifier (2)<br>- Molecular weight (2)<br>- Default structural alerts + unwanted moiety (1) |

**Supplementary Table 9.** Number of high-scoring generated molecules, total analogs and high scoring-analogs found in catalogs and final number of molecules tested. *Up to 10 neighbors per generated molecule. **We tested only the closest high-scoring molecule to each generated compound.

| Exercise | # high scoring generated molecules | # similar molecules found in DBs* | # high scoring molecules found in DBs | # molecules tested** |
| --- | --- | --- | --- | --- |
| BXPC3 vs CAPAN2 | 137 | 1,154 | 105 | 16 |
| BXPC3 vs HEK293 | 47 | 445 | 93 | 22 |
| CAPAN2 vs HEK293 | 100 | 87 | 1 | 1 |
| CAPAN2 vs CFPAC1 | 121 | 167 | 5 | 1 |
| HEK293 vs BXPC3 | 60 | 415 | 33 | 1 |
| HEK293 vs CAPAN2 | 37 | 267 | 7 | 1 |
| HEK293 vs CFPAC1 | 37 | 74 | 6 | 11 |

**Supplementary Table 10.** Experimental validation of generated molecules vs IRB library predictions. The table reports initial validations at 1 and 10 µM and subsequent dose–response curves for the identified hits.

**Supplementary Table 11.** Table with abbreviation and meaning of the 11 Bioteque metapaths used as cell line descriptors to compare to cell line sensitivity (Extended Data Fig. 3). *Dataset names as they appear in the Bioteque^25^ web resource, where we downloaded the descriptors from.

| Metapath abbreviation | Metapath full name | Dataset* |
| --- | --- | --- |
| CLL-ass-PWY | Cell associates Pathway | Cellosaurus-Cosmic |
| CLL-cnd-GEN | Cell copy number down Gene | GDSC |
| CLL-cnu-GEN | Cell copy number up Gene | CCLE_hmz |
| CLL-dwr-GEN | Cell downregulates Gene | GDSC |
| CLL-dwr-GEN-ass-PWY | Cell downregulates Gene (that) associates Pathway | GDSC - Reactome |
| CLL-dwr-GEN-has-MFN | Cell downregulates Gene (that) has Molecular Function | GDSC - GO |
| CLL-dwr-GEN-ppi-GEN | Cell downregulates Gene (that) protein-protein interacts Gene | GDSC - STRING |
| CLL-dwr+upr-GEN-dwr+upr-CLL | Cell downregulates or upregulates Gene | GDSC - GDSC |
| CLL-mth-GEN | Cell methylates Gene | GDSC |
| CLL-mut-GEN | Cell mutation Gene | CCLE |
| CLL-mut-GEN-ass-PWY | Cell mutation Gene (that) associates Pathway | GDSC - Reactome |
| CLL-sns-CPD | Cell sensitive Compound | Drugcell |
| CLL-upr-GEN | Cell upregulates Gene | GDSC |
| CLL-upr-GEN-ass-PWY | Cell upregulates Gene (that) associates Pathway | GDSC - Reactome |

## Source data

All the experimental data generated in this study is publicly available. See *Data Availability*.

## Notes

https://github.com/sbnb-irb/cytotoxics_design.git

### Competing Interest Statement

The authors have declared no competing interest.

## References

1. Macarron, R. et al. Impact of high-throughput screening in biomedical research. Nat. Rev. Drug Discov. 10, 188–195 (2011).

2. Warr, W. A., Nicklaus, M. C., Nicolaou, C. A. & Rarey, M. Exploration of Ultralarge Compound Collections for Drug Discovery. J. Chem. Inf. Model. 62, 2021–2034 (2022).

3. Lyu, J., Irwin, J. J. & Shoichet, B. K. Modeling the expansion of virtual screening libraries. Nat. Chem. Biol. 19, 712–718 (2023).

4. Gentile, F. et al. Artificial intelligence-enabled virtual screening of ultra-large chemical libraries with deep docking. Nat. Protoc. 17, 672–697 (2022).

5. Sivula, T. et al. Machine Learning-Boosted Docking Enables the Efficient Structure-Based Virtual Screening of Giga-Scale Enumerated Chemical Libraries. J. Chem. Inf. Model. 63, 5773–5783 (2023).

6. Cao, Z., Sciabola, S. & Wang, Y. Large-Scale Pretraining Improves Sample Efficiency of Active Learning-Based Virtual Screening. J. Chem. Inf. Model. 64, 1882–1891 (2024).

7. Serrano-Morrás, Á. et al. A bottom-up approach to find lead compounds in expansive chemical spaces. Commun. Chem. 8, 225 (2025).

8. Zhavoronkov, A. et al. Deep learning enables rapid identification of potent DDR1 kinase inhibitors. Nat. Biotechnol. 37, 1038–1040 (2019).

9. Isigkeit, L. et al. Automated design of multi-target ligands by generative deep learning. Nat. Commun. 15, 7946 (2024).

10. Blaschke, T. et al. REINVENT 2.0: An AI Tool for De Novo Drug Design. J. Chem. Inf. Model. 60, 5918– 5922 (2020).

11. Loeffler, H. H. et al. Reinvent 4: Modern AI–driven generative molecule design. J. Cheminformatics 16, 20 (2024).

12. Du, Y. et al. Machine learning-aided generative molecular design. *Nat*. Mach. Intell. 6, 589–604 (2024).

13. Zhung, W., Kim, H. & Kim, W. Y. 3D molecular generative framework for interaction-guided drug design. Nat. Commun. 15, 2688 (2024).

14. Behan, F. M. et al. Prioritization of cancer therapeutic targets using CRISPR–Cas9 screens. Nature 568, 511–516 (2019).

15. Tsherniak, A. et al. Defining a Cancer Dependency Map. Cell 170, 564–576.e16 (2017).

16. Sadri, A. Is Target-Based Drug Discovery Efficient? Discovery and “Off-Target” Mechanisms of All Drugs. J. Med. Chem. 66, 12651–12677 (2023).

17. Swinney, D. C. & Anthony, J. How were new medicines discovered? Nat. Rev. Drug Discov. 10, 507– 519 (2011).

18. Parisi, D. et al. Drug repositioning or target repositioning: A structural perspective of drug-target-indication relationship for available repurposed drugs. Comput. Struct. Biotechnol. J. 18, 1043–1055 (2020).

19. Kawata, K. et al. Trans-omic Analysis Reveals Selective Responses to Induced and Basal Insulin across Signaling, Transcriptional, and Metabolic Networks. iScience 7, 212–229 (2018).

20. Vitrinel, B. et al. Exploiting Interdata Relationships in Next-generation Proteomics Analysis. Mol. Cell. Proteomics MCP 18, S5–S14 (2019).

21. Subramanian, A. et al. A Next Generation Connectivity Map: L1000 Platform and the First 1,000,000 Profiles. Cell 171, 1437–1452.e17 (2017).

22. Mitchell, D. C. et al. A proteome-wide atlas of drug mechanism of action. Nat. Biotechnol. 41, 845–857 (2023).

23. Dubuis, S., Ortmayr, K. & Zampieri, M. A framework for large-scale metabolome drug profiling links coenzyme A metabolism to the toxicity of anti-cancer drug dichloroacetate. *Commun*. Biol. 1, 101 (2018).

24. Diamant, I., Clarke, D. J. B., Evangelista, J. E., Lingam, N. & Ma’ayan, A. Harmonizome 3.0: integrated knowledge about genes and proteins from diverse multi-omics resources. Nucleic Acids Res. 53, D1016–D1028 (2025).

25. Fernández-Torras, A., Duran-Frigola, M., Bertoni, M., Locatelli, M. & Aloy, P. Integrating and formatting biomedical data as pre-calculated knowledge graph embeddings in the Bioteque. Nat. Commun. 13, 5304 (2022).

26. Fernández-Torras, A., Locatelli, M., Bertoni, M. & Aloy, P. BQsupports: systematic assessment of the support and novelty of new biomedical associations. Bioinformatics 39, btad581 (2023).

27. Duran-Frigola, M. et al. Extending the small-molecule similarity principle to all levels of biology with the Chemical Checker. Nat. Biotechnol. 38, 1087–1096 (2020).

28. Bertoni, M. et al. Bioactivity descriptors for uncharacterized chemical compounds. Nat. Commun. 12, 3932 (2021).

29. Comajuncosa-Creus, A. et al. Integration of diverse bioactivity data into the Chemical Checker compound universe. Preprint at 10.1101/2024.12.04.626832 (2024).

30. Joo, S., Kim, M. S., Yang, J. & Park, J. Generative Model for Proposing Drug Candidates Satisfying Anticancer Properties Using a Conditional Variational Autoencoder. ACS Omega 5, 18642–18650 (2020).

31. Méndez-Lucio, O., Baillif, B., Clevert, D.-A., Rouquié, D. & Wichard, J. De novo generation of hit-like molecules from gene expression signatures using artificial intelligence. Nat. Commun. 11, 10 (2020).

32. Matsukiyo, Y., Tengeiji, A., Li, C. & Yamanishi, Y. Transcriptionally Conditional Recurrent Neural Network for De Novo Drug Design. J. Chem. Inf. Model. 64, 5844–5852 (2024).

33. Yamanaka, C., Uki, S., Kaitoh, K., Iwata, M. & Yamanishi, Y. De novo drug design based on patient gene expression profiles via deep learning. Mol. Inform. 42, e2300064 (2023).

34. Liu, H., Tian, S. & Liu, X. Phenotypic Profile-Informed Generation of Drug-Like Molecules via Dual-Channel Variational Autoencoders. Preprint at 10.48550/arXiv.2506.02051 (2025).

35. Kim, H. et al. A genotype-to-drug diffusion model for generation of tailored anti-cancer small molecules. Nat. Commun. 16, 5628 (2025).

36. Shoemaker, R. H. The NCI60 human tumour cell line anticancer drug screen. Nat. Rev. Cancer 6, 813– 823 (2006).

37. Rees, M. G. et al. Correlating chemical sensitivity and basal gene expression reveals mechanism of action. Nat. Chem. Biol. 12, 109–116 (2016).

38. Yang, W. et al. Genomics of Drug Sensitivity in Cancer (GDSC): a resource for therapeutic biomarker discovery in cancer cells. Nucleic Acids Res. 41, D955–961 (2013).

39. Kuenzi, B. M. et al. Predicting Drug Response and Synergy Using a Deep Learning Model of Human Cancer Cells. Cancer Cell 38, 672–684.e6 (2020).

40. Safikhani, Z. et al. Revisiting inconsistency in large pharmacogenomic studies. F1000Research 5, 2333 (2016).

41. Pozdeyev, N. et al. Integrating heterogeneous drug sensitivity data from cancer pharmacogenomic studies. Oncotarget 7, 51619–51625 (2016).

42. DepMap, B. DepMap 24Q4 Public. 30825074613 Bytes Figshare+ 10.25452/FIGSHARE.PLUS.27993248.V1 (2024).

43. Rogers, D. & Hahn, M. Extended-Connectivity Fingerprints. J. Chem. Inf. Model. 50, 742–754 (2010).

44. Irwin, J. J. et al. ZINC20—A Free Ultralarge-Scale Chemical Database for Ligand Discovery. J. Chem. Inf. Model. 60, 6065–6073 (2020).

45. Kiss, R., Sandor, M. & Szalai, F. A. http://Mcule.com: a public web service for drug discovery. J. Cheminformatics 4, P17, 1758-2946-4-S1-P17 (2012).

46. Grygorenko, O. O. et al. Generating Multibillion Chemical Space of Readily Accessible Screening Compounds. iScience 23, 101681 (2020).

47. SmallWorld | Load. https://sw.docking.org/api.html.

48. Murphy, K. M. et al. Evaluation of candidate genes MAP2K4, MADH4, ACVR1B, and BRCA2 in familial pancreatic cancer: deleterious BRCA2 mutations in 17%. Cancer Res. 62, 3789–3793 (2002).

49. Olaoba, O. T. et al. Driver Mutations in Pancreatic Cancer and Opportunities for Targeted Therapy. Cancers 16, 1808 (2024).

50. Zhang, Q. et al. Fbxw7 Deletion Accelerates KrasG12D-Driven Pancreatic Tumorigenesis via Yap Accumulation. Neoplasia 18, 666–673 (2016).

51. Barretina, J. et al. The Cancer Cell Line Encyclopedia enables predictive modelling of anticancer drug sensitivity. Nature 483, 603–607 (2012).

52. The Cancer Cell Line Encyclopedia Consortium & The Genomics of Drug Sensitivity in Cancer Consortium. Pharmacogenomic agreement between two cancer cell line data sets. Nature 528, 84–87 (2015).

53. Smirnov, P. et al. PharmacoGx: an R package for analysis of large pharmacogenomic datasets. Bioinformatics 32, 1244–1246 (2016).

54. Krueger, F. et al. Trim Galore v0.6.10. Zenodo 10.5281/zenodo.7598955 (2023).

55. Quast, C. et al. The SILVA ribosomal RNA gene database project: improved data processing and web-based tools. Nucleic Acids Res. 41, D590–D596 (2012).

56. Langmead, B. & Salzberg, S. L. Fast gapped-read alignment with Bowtie 2. Nat. Methods 9, 357–359 (2012).

57. Dobin, A. et al. STAR: ultrafast universal RNA-seq aligner. Bioinforma. Oxf. Engl. 29, 15–21 (2013).

58. Liao, Y., Smyth, G. K. & Shi, W. The Subread aligner: fast, accurate and scalable read mapping by seed-and-vote. Nucleic Acids Res. 41, e108–e108 (2013).

59. Iorio, F. et al. A Landscape of Pharmacogenomic Interactions in Cancer. Cell 166, 740–754 (2016).

